# Stabilization of AMPK/PFKL/RPIA in the Glycolytic Bodies Transduces IL6/STAT3 Signal in Hepatocarcinogenesis

**DOI:** 10.1101/2024.02.29.582877

**Authors:** He-Yun Hsiao, Chun-Chia Cheng, Yu-Ting Chou, Cheng-Chin Kuo, Wen-Ching Wang, Bonifasius Putera Sampurna, Yi-Wen Wang, Chun-Ling Hsiao, Jing-Yiing Wu, Kuan-Hao Lin, Wan-Yu Yang, Yu-Hsuan Lin, Kong-Huai Gwee, Horng-Dar Wang, Chiou-Hwa Yuh

**Affiliations:** Institute of Molecular and Genomic Medicine, National Health Research Institutes, Taiwan; Institute of Biotechnology, National Tsing Hua University, Hsinchu, Taiwan; Radiation Biology Research Center, Institute for Radiological Research, Chang Gung University/Chang Gung Memorial Hospital, Linkou, Taiwan; Department of Medicine, Division of Hematology and Oncology, and the Helen Diller Comprehensive Cancer Center, University of California, San Francisco, USA; Institute of Cellular and Systems Medicine, National Health Research Institutes, Taiwan; Institute of Molecular and Cellular Biology, National Tsing Hua University, Hsinchu, Taiwan; Institute of Bioinformatics and Structural Biology, National Tsing Hua University, Hsinchu, Taiwan; Department of Biological Science and Technology, National Yang Ming Chiao Tung University, Hsinchu, Taiwan; Ph.D. Program in Environmental and Occupational Medicine, Kaohsiung Medical University, Kaohsiung, Taiwan

**Keywords:** PFKL, RPIA, AMPK, IL6, STAT3, glycolytic body, liver cancer, metastasis

## Abstract

Metabolic reprogramming is a pivotal characteristic of cancer, yet the intricate interplay between glycolysis and the pentose phosphate pathway (PPP) remains elusive. This study unveils the pivotal role of 6-phosphofructokinase liver type (PFKL) in glycolysis and ribose 5-phosphate isomerase A (RPIA) in PPP, orchestrating liver tumorigenesis. PFKL, the rate-limiting enzyme in glycolysis, stabilizes RPIA by impeding ubiquitination/proteasome activity. The pro-inflammatory and tumor cytokine interleukin 6 activates pSTAT3 which binds to the promoter region and activates AMPK and PFKL transcription. Furthermore, pAMPK stabilizes PFKL protein by preventing proteasome degradation in hepatoma cells. Inhibiting PFKL, AMPK, and STAT3 genetically or pharmacologically can reduce glycolysis, ATP production, resulting in reduction of hepatoma cell proliferation and migration. Intriguingly, the PFKL, AMPK, RPIA, and PKM2 are co-localized in the Glycolytic body (G-body) which starts forming at chronic hepatitis, dramatically increases during active hepatitis, and the size of G-bodies becomes bigger from cirrhosis to hepatocellular carcinoma. Furthermore, using Bimolecular fluorescence complementation (BiFC) assay, we demonstrated that PFKL and RPIA direct interacts. Targeting AMPK or STAT3 significantly reduced tumor formation and lipid accumulation in zebrafish models, suggesting the STAT3/AMPK/PFKL axis as a potential therapeutic avenue for liver cancer treatment.

## Introduction

Liver cancer, specifically hepatocellular carcinoma (HCC), is the fourth leading cause of cancer-related fatalities and driven by factors such as hepatitis infections, aflatoxin exposure, alcohol abuse, and Metabolic Dysfunction-associated fatty liver disease (MALFD)/Nonalcoholic fatty liver disease (NAFLD) [1, 2]. Treatment options are diverse, ranging from surgery and transplantation to therapies such as ablation, chemoembolization, and immunotherapy, varying based on cancer stage and overall health [3]. Advanced HCC management includes approved drugs such as sorafenib and lenvatinib, categorized as tyrosine kinase inhibitors, along with combinations such as atezolizumab and bevacizumab, blending immune checkpoint inhibitors with antiangiogenic agents, and offering multiple treatment routes [3, 4]. However, patients with HCC associated with nonalcoholic steatohepatitis (NASH) often exhibit poorer responses, particularly to immunotherapies, due to compromised immunity [5]. Despite the promising nature of targeted therapies, their efficacy benefits only a limited patient subset, underscoring the imperative for more efficacious strategies, especially for HCC associated with NAFLD [6]. The complexities posed by HCC, especially in cases induced by NASH, necessitate urgent advancements in therapeutic strategies to improve patient outcomes.

Cancer is characterized by a profound metabolic shift, and one pivotal aspect of this alteration is the phenomenon of metabolic reprogramming. Phosphofructokinase-1 (PFK1) is a critical enzyme in glycolysis, catalyzing the conversion of fructose-6-phosphate into fructose-1,6-bisphosphate [7]. Intriguingly, the upregulation of liver-type PFK1, denoted PFKL, has been correlated with unfavorable survival outcomes in lung cancer [8]. The significance of PFKL lies in its role in meeting the heightened glucose demands of tumor cells, rendering it a prospective target for anticancer interventions [9, 10]. IL6 secreted by activated Kupffer cells within the inflamed liver, plays a pivotal role in HCC development [11]. IL6 exerts its influence by phosphorylating signal transducer and activator of transcription 3 (STAT3), a transcription factor that is a crucial regulator of HCC tumorigenesis [12–14]. This IL6/STAT3-mediated tumorigenesis has been extensively studied and is intrinsically linked to glycolysis, notably through its regulation of 6-phosphofructo-2-kinase/fructose-2,6-biphosphatase 3 (PFKFB3) in colorectal cancer [15]. Moreover, another downstream key enzyme of PFKFB3 in glycolysis is PFK1 [16], which goes beyond determining the glycolytic rate [17]. Remarkably, PFK1 assembles into nonmembrane-bound glycolytic bodies, referred to as G-bodies, within liver cancer HepG2 cells under hypoxic conditions [18]. These findings underscore the intricate association between IL6/STAT3 signaling and aerobic glycolysis in tumorigenesis.

Accumulating evidence emphasizes the critical role of dysregulated metabolism as a primary driver of carcinogenesis, with the glycolytic phenotype notably associated with enhanced cell proliferation and metastasis [19]. Tumor cells often employ the Warburg effect as a metabolic strategy to secure the requisite energy for their accelerated growth and metastatic potential. The intricate coordination of glycolysis, the PPP pathway, and lipid metabolism plays a paramount role in hepatocarcinogenesis. Our previous research revealed that the upregulation of RPIA, a pivotal enzyme within the PPP pathway, serves as a catalyst for hepatocarcinogenesis through the activation of extracellular signal-regulated kinase (ERK) [20]. In the zebrafish model, the overexpression of RPIA not only elevated the levels of phosphorylated AMP-activated protein kinase (pAMPK) and phosphorylated ERK during steatosis [21] but also led to heightened pERK and β-catenin activity during cancer formation [21]. Extended from our above studies, we delve into the role of PFKL in HCC and further explore the intricate interplay between PFKL and RPIA in this context.

IL6 has also been shown to enhance AMPK phosphorylation in muscle tissues [22, 23], a response typically activated during energy deprivation. While the role of AMPK in tumorigenesis remains debatable, some current literature suggests that AMPK acts as a tumor suppressor by downregulating target of rapamycin (mTOR) [24, 25], yet there is also evidence indicating that AMPK activation can trigger tumorigenesis [24, 26]. Notably, Snf1p, a yeast homolog of AMPK, modulates G-body formation [18], supporting the notion that AMPK may play a role in PFKL-mediated glycolysis in liver cancer. Furthermore, other research has highlighted targeting AMPK to suppress tumor initiation and progression [24]. This is exemplified by metformin, which predominantly inhibits mTOR, leading to increased AMP and ADP levels and consequently activating AMPK [24]. It has been proposed that combining metformin with an AMPK inhibitor could enhance therapeutic efficacy in cancer [24]. In this study, we conducted an in-depth exploration of the regulatory mechanisms governing the expression of PFKL and RPIA in liver cancer cells. We performed immunohistochemistry and immunofluorescence staining for the co-localization of PFKL, RPIA, AMPK and PKM2 in human diseases specimens. Additionally, we investigated the inhibitory effects of targeting STAT3 and AMPK on hepatoma cell proliferation using xenotransplantation in zebrafish. Our research evaluated the impact of these interventions on reducing lipid accumulation and preventing liver cancer formation in transgenic zebrafish models. Moreover, our research sheds light on the clinical relevance of AMPK/PFKL/RPIA in G-bodies in hepatocarcinogenesis.

## Results

### PFKL and RPIA collaborate in promoting hepatoma cell proliferation

To examine if PFKL plays a role in liver cancer formation, we utilized a tissue array encompassing all stages of HCC specimens for IHC analysis to assess PFKL expression levels. Elevated PFKL protein levels were detected in various stages of liver cancer, with an initial increase observed as early as stage I and intensifying through advanced stages (**Fig. 1A**). Quantification of immunoreactivity scores (IRS) revealed a significant increase in PFKL expression in HCC compared to nonneoplastic liver tissue, with scores rising across different advanced HCC stages, except for a slight drop in the metastatic stage (**Fig. 1A**). These findings highlight the upregulation of PFKL during HCC progression.

**Figure 1.**
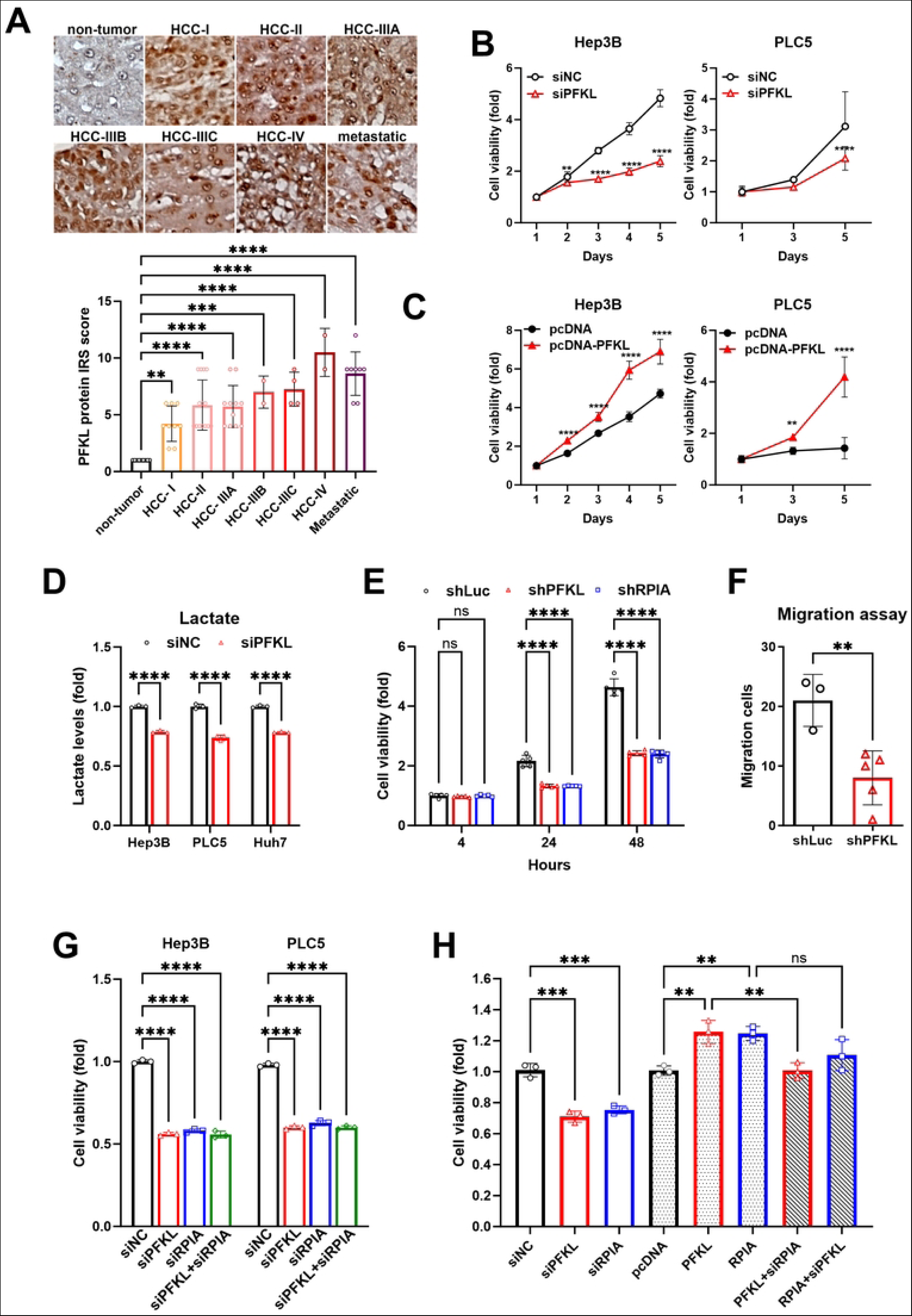
Coordinated actions of PFKL and RPIA in hepatocellular carcinoma (HCC). (**A**) Immunostaining of PFKL in HCC tissues compared to nonneoplastic liver tissues was carried out, followed by quantification using the immunoreactive score (IRS) method. Staining intensity (1 to 3) is combined with the percentage of positively stained cells (1 to 4). (**B**) Knockdown of PFKL reduces viability in Hep3B and PLC5 hepatoma cells. The red line indicates siPFKL, while the black line denotes the siNC control. (**C**) Overexpression of PFKL increases viability in Hep3B and PLC5 hepatoma cells. The red line indicates pcDNA-PFKL, and the black line denotes pcDNA control. (**D**) Knockdown of PFKL significantly reduces lactate production in hepatoma cells. The red bar indicates siPFKL, and the black bar denotes the siNC control. (**E**) Knockdown of either PFKL or RPIA with shRNA reduces viability in PLC5 hepatoma cells. The red bar indicates shPFKL, the blue bar denotes shRPIA, and the black bar represents shLuc control. (**F**) Knockdown of PFKL reduces the migration ability of PLC5 cells. The red bar indicates shPFKL, and the black bar denotes shLuc control. (**G**) Simultaneous knockdown of PFKL and RPIA does not result in any synergistic reduction in proliferation in Hep3B and PLC5 cells. The black bar indicates shLuc control, the red bar denotes shPFKL, the blue bar represents shRPIA, and the green bar represents the combination of siPFKL and siRPIA. (**H**) PFKL overexpression-induced proliferation in hepatoma cells can be reversed by RPIA knockdown. The red bar indicates siPFKL, the blue bar denotes siRPIA, and the black bar represents the siNC control. Bars with light gray coloring inside indicate pcDNA control, PFKL, and RPIA, while the bars with dark gray coloring inside represent PFKL+siRPIA or RPIA+siPFKL. Statistical analyses were performed using one-way ANOVA. **p < 0.01; ***p < 0.001; ****p < 0.0001.

To investigate the effect of PFKL expression on liver cancer proliferation, we conducted knockdown and overexpression of PFKL in hepatoma cells. Knockdown of PFKL significantly reduced cell viability in Hep3B and PLC5 hepatoma cells (**Fig. 1B**), while PFKL overexpression exhibited the opposite effect (**Fig. 1C**). An issue arises when perturbations affect glycolytic metabolism, potentially leading to alterations in WST-1 reduction independent of changes in cell viability. To eliminate this concern, we employed fluorescence-activated cell sorting (FACS) to validate any cell cycle alterations resulting from PFKL knockdown or overexpression (**Fig. S1**), and the results support our findings. Given that lactate production is typically associated with glycolysis rates in cancer cells, we also observed a reduction in lactate production in hepatoma cells upon PFKL knockdown (**Fig. 1D**). These results reinforce the role of PFKL in enhancing liver cancer cell proliferation, supporting our previous observations of PFKL upregulation in HCC formation.

Building upon our prior findings that RPIA regulates cell proliferation in liver cancer cells [20], we detected that knockdown of either PFKL or RPIA similarly reduced cell viability (**Fig. 1E**), and PFKL knockdown additionally decreased migration activity in PLC5 cells (**Fig. 1F**). This prompted us to investigate the potential interplay between PFKL and RPIA in regulating cancer cell proliferation. Intriguingly, simultaneous knockdown of PFKL and RPIA did not result in a further reduction in cell viability compared to knockdown of each individual gene in Hep3B and PLC5 cells (**Fig. 1G**), suggesting that both genes may function within the same pathway to modulate liver cancer cell proliferation. Moreover, PFKL overexpression-mediated increased cell proliferation could be fully suppressed by RPIA knockdown, yet RPIA overexpression-induced cell proliferation was not significantly reduced by PFKL knockdown (**Fig. 1H**), implying a potential epistatic relationship between PFKL and RPIA in liver cancer cell proliferation. These results suggest that PFKL and RPIA operate within the same axis to regulate liver cancer cell proliferation.

### PFKL stabilizes RPIA protein levels via the ubiquitination-proteasome pathway

To further explore the relationship between PFKL and RPIA, we conducted knockdown experiments for both PFKL and RPIA and assessed their respective protein levels in liver cancer cells. Interestingly, we observed that RPIA protein levels decreased significantly upon PFKL knockdown; conversely, RPIA knockdown had no discernible impact on PFKL protein levels (**Fig. 2A**). Intriguingly, PFKL knockdown did not significantly reduce RPIA mRNA expression in the PLC5 cell line but did so to a small extent in the Hep3B cell line (**Fig. 2B**). These results indicate that PFKL regulates RPIA at the protein level independently of transcriptional regulation.

**Figure 2.**
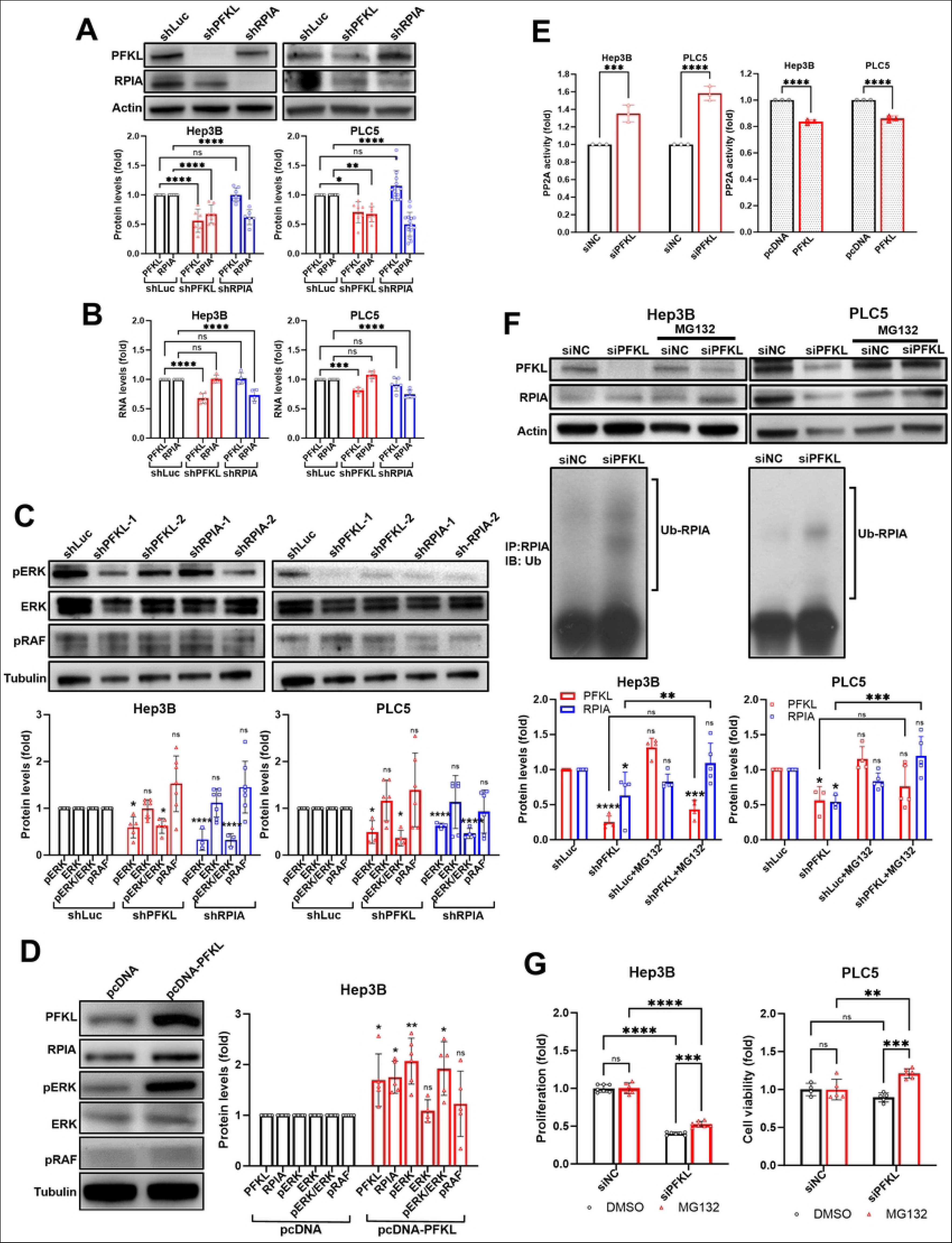
PFKL modulates RPIA stability via the ubiquitination/proteasome pathway. (**A**) Knockdown of PFKL reduces RPIA protein levels, but knockdown of RPIA does not affect PFKL protein levels in hepatoma cell lines. The lower panel depicts the quantification results of western blot analysis for PFKL and RPIA protein levels. The black bar represents the shLuc control, the red bar signifies shPFKL, and the blue bar represents shRPIA. (**B**) Knockdown of PFKL does not affect RPIA mRNA expression, and knockdown of RPIA does not change PFKL mRNA levels in hepatoma cells as determined by quantitative PCR (qPCR), suggesting that PFKL knockdown-mediated RPIA protein reduction does not occur at the transcriptional level. The black bar indicates shLuc control, the red bar denotes shPFKL, and the blue bar represents shRPIA. (**C**) Knockdown of PFKL decreases pERK protein levels but does not affect pRaf. The lower panel presents quantification results of western blot analysis for pERK, total ERK, the ratio of pERK/ERK, and pRAF. The black bar represents shLuc control, the red bar signifies shPFKL, and the blue bar represents shRPIA. (**D**) Overexpression of PFKL increases PFKL, RPIA, and pERK protein levels but has no effect on pRaf. The right panel illustrates the quantification results of western blot analysis for PFKL, RPIA, pERK, total ERK, the ratio of pERK/ERK, and pRAF. The black bar indicates shLuc control, the red bar denotes shPFKL, and the blue bar represents shRPIA. (**E**) Knockdown of PFKL increases PP2A activity, while overexpression of PFKL reduces PP2A activity. Quantitative results for PP2A activity are shown for PFKL knockdown (red bar) versus siNC control (black bar) in Hep3B and PLC5 cells and for PFKL overexpression (red bar with dots inside) versus pcDNA control (black bar with dots inside). (**F**) MG132 (protease inhibitor) treatment rescues the PFKL knockdown-mediated downregulation of RPIA in both hepatoma cell lines (upper panel). Knockdown of PFKL elevates ubiquitinated RPIA levels in both hepatoma cell lines (middle panel). The lower panel displays quantification results for PFKL protein (red bar) and RPIA protein (blue bar) in shLuc control, shPFKL, MG132, and shPFKL+MG132 cotreatment. (**G**) MG132 rescues the PFKL knockdown-induced suppression of cell viability. The red bar indicates MG132 treatment, and the black bar denotes the DMSO control in siNC versus siPFKL. Statistical analyses were conducted using one-way ANOVA. **p < 0.01; ***p < 0.001; ****p < 0.0001.

Previously, we found that RPIA induces phosphorylated extracellular signal-regulated kinas (pERK) in hepatoma cells [20] and in a zebrafish model [21]. In this study, we found that PFKL knockdown not only reduced RPIA levels but also decreased pERK levels yet had no effect on pRaf (**Fig. 2C**), while PFKL overexpression again showed the opposite effect (**Fig. 2D**). Importantly, this interrelationship between PFKL and RPIA appeared to be specific, as knockdown of both genes did not affect the expression of other glycolytic enzymes, such as pyruvate kinase (PK), phosphoglucose isomerase (PGI), and triose-phosphate isomerase (TPI) (**Fig. S2A**), nor did it affect other enzymes in the PPP, such as glucose-6-phosphate dehydrogenase (G6PD), transaldolase (TALDO), and transketolase (TKL) (**Fig. S2B**). PFKL specifically increased RPIA and cMyc oncoprotein levels in liver cancer cells (**Fig. S2C**).

Previously, we demonstrated that RPIA overexpression led to elevated pERK levels, attributed to decreased PP2A phosphatase activity in hepatoma cells [20]. If PFKL can function upstream of RPIA, we asked whether PFKL can modulate PP2A activity similar to RPIA. The results indicated that PFKL knockdown increased PP2A activity, while PFKL overexpression reduced PP2A activity in hepatoma cell lines (**Fig. 2E**). These findings further support the notion that PFKL may operate within the same signaling pathway as RPIA.

Since PFKL regulates RPIA at the protein level, we next hypothesized that PFKL knockdown may trigger the degradation of RPIA through the ubiquitin‒proteasome pathway. We employed an MG132 protease inhibitor to test this hypothesis and found that MG132 treatment rescued the PFKL knockdown-induced reduction in RPIA protein levels (**Fig. 2F**). Furthermore, PFKL knockdown increased the ubiquitination of RPIA (**Fig. 2F**). Moreover, MG132 reversed the inhibitory effect of siPFKL on hepatoma cell viability (**Fig. 2G**). These results suggest that PFKL regulates RPIA post-translationally by preventing the ubiquitin‒proteasome degradation of RPIA in hepatoma cells, highlighting the specific interaction between PFKL in glycolysis and RPIA in the pentose phosphate pathway.

### IL6 induces PFKL expression via AMPK, stabilizing the PFKL protein and protecting it from proteasomal degradation

Considering the pivotal role of IL6 in liver tumorigenesis and its influence on glycolysis, we examined whether IL6 could upregulate PFKL and RPIA mRNA levels in PLC5 cells by qPCR analysis. The results unequivocally revealed that IL6 substantially elevated the mRNA levels of critical glycolysis-related genes, such as Glut1, HK2, PFKFB3, PFKL, PKM2, LDHA, and RPIA, while having no discernible impact on RPE in the PPP (**Fig. 3A**). Furthermore, we utilized cancer sphere formation to isolate and study cancer stem cells (CSCs). Interestingly, we observed even more pronounced elevations in the mRNA levels of these glycolytic and PPP pathway genes (**Fig. 3A**). Furthermore, IL6 treatment significantly induced the protein levels of PFKL in PLC5 cells (**Fig. 3B**).

**Figure 3.**
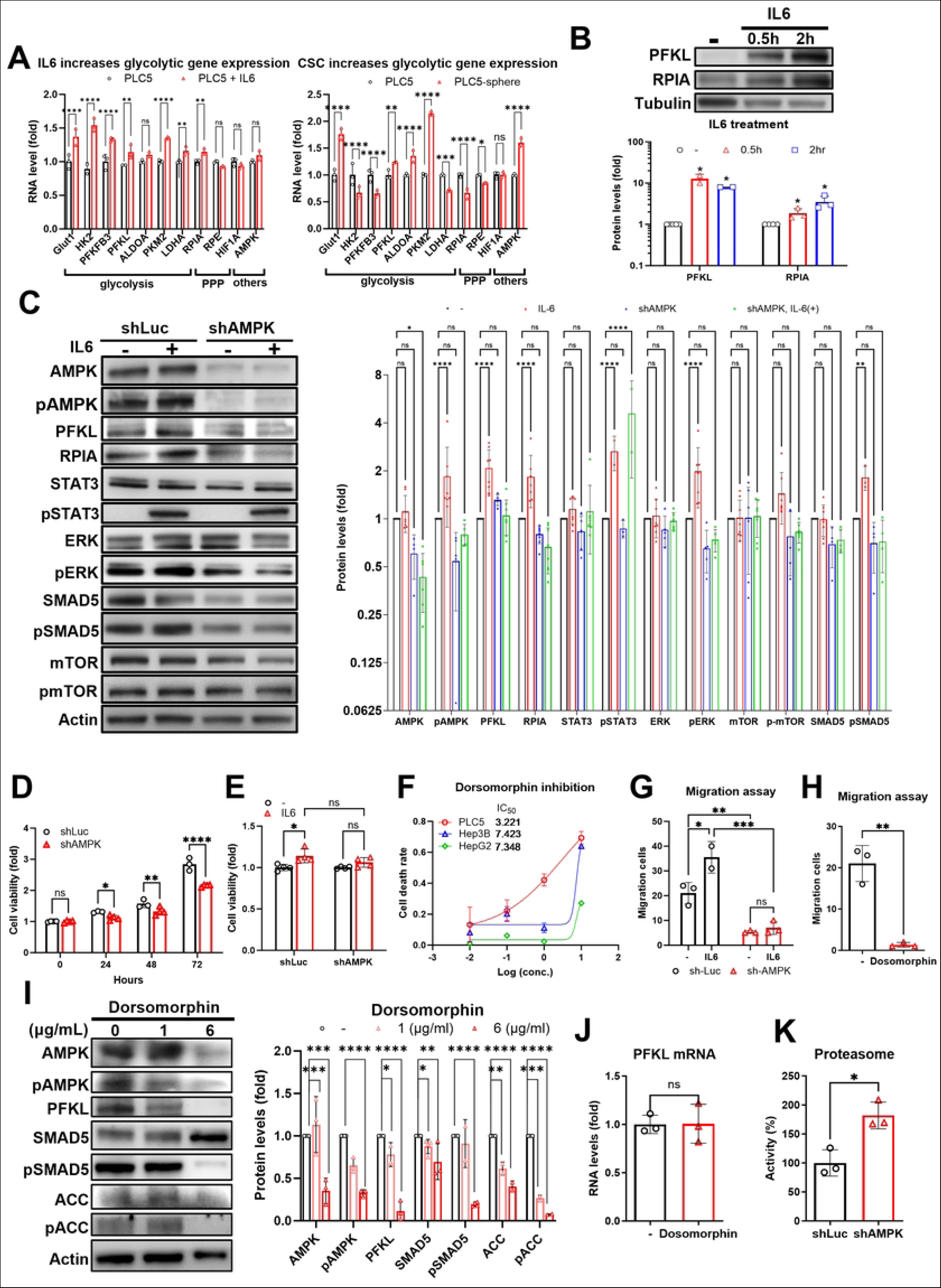
AMPK is needed in IL6-induced PFKL-upregulated protein levels in PLC5 cells. (**A**) Treatment with IL6 (20 ng/mL) for 2 hours (left panel) or the induction of cancer stem cells (CSCs) (right panel) upregulates the expression of most glycolysis-related genes, including PFKL, at the mRNA level. Red bars represent PLC5 cells after IL6 treatment or sphere formation, while black bars represent control cells. (**B**) IL6 treatment increases PFKL protein levels. The right panel shows the quantification results of western blot analysis for PFKL and RPIA protein levels. Black bars indicate no treatment control, red bars represent 0.5 hours of IL6 treatment, and blue bars represent 2 hours of IL6 treatment. (**C**) Knockdown of AMPK by shRNA reduces IL6-induced PFKL-upregulated protein expression in PLC5 cells, leading to decreases in pAMPK, PFKL, RPIA, pERK, and pSMAD5 levels, while it has no effect on STAT3 phosphorylation. The right panel illustrates the quantification of western blot results for AMPK, pAMPK, PFKL, RPIA, STAT3, pSTAT3, ERK, pERK, mTOR, p-mTOR, SMAD5, and pSMAD5. Black bars represent no treatment control, red bars indicate IL6 treatment, blue bars represent shAMPK, and the green bar denotes shAMPK+IL6. (**D**) Knockdown of AMPK decreases cell viability in PLC5 cells. Quantification of cell viability at 24, 48, and 72 hours is shown. Red bars represent shAMPK, while black bars denote shLuc control. (**E**) AMPK knockdown reduces IL6-stimulated cell viability, normalized to the control without IL6 treatment. Black bars represent the control without treatment, and red bars represent IL6 treatment. (**F**) Inhibition of AMPK with dorsomorphin decreases cell viability in three hepatoma cell lines. Cell death rates were quantified, and the IC50 for dorsomorphin in PLC5, Hep3B, and HepG2 cells is displayed in the upper left. (**G**) AMPK knockdown decreases migration ability with or without IL6 treatment. Quantification of migration without IL6 or with IL6 treatment is shown. Black bars represent sh-Luc control, and red bars denote sh-AMPK. (**H**) Suppression of AMPK with dorsomorphin reduces migration ability. Black bars represent the control without treatment, and red bars indicate dorsomorphin treatment. (**I**) Dorsomorphin significantly reduces PFKL protein levels. The left panel presents quantification of western blot results for AMPK, pAMPK, PFKL, SMAD5, pSMAD5, ACC, and pACC. Black bars represent the control without treatment, light red bars denote 1 µg/mL, and red bars indicate 6 µg/mL dorsomorphin treatment. (**J**) AMPK knockdown does not affect PFKL mRNA expression. Quantification of qPCR results for PFKL mRNA is shown. The red bar indicates dorsomorphin treatment, while the black bar denotes the control without treatment. (**K**) AMPK knockdown increases proteasome activity in PLC5 cells, suggesting that AMPK stabilizes PFKL by inhibiting proteasome activity. Quantification of proteasome activity is shown, with the red bar indicating shAMPK and the black bar representing shLuc control. Statistical analyses were performed using one-way ANOVA. *p < 0.05; **p < 0.01; ***p < 0.001; ****p < 0.0001.

To gain insights into how IL6 augments PFKL expression, we conducted AMPK knockdown experiments since AMPK is one of the downstream effectors of IL6. We measured protein levels in the presence or absence of IL6 treatment. Our results indicated that IL6 elevated the levels of pAMPK, PFKL, RPIA, pSTAT3, pERK, and pSMAD5, and this effect was attenuated by AMPK knockdown but did not significantly affect the levels of pSTAT3 (**Fig. 3C**). This suggests that IL6 enhances the expression of PFKL, RPIA, and ERK through AMPK, with pSTAT3 potentially acting upstream of AMPK activation. Knockdown of AMPK resulted in reduced cancer cell viability (**Fig. 3D**). Conversely, the IL6-mediated increase in cancer cell viability could be blocked by AMPK knockdown (**Fig. 3E**). To corroborate the role of AMPK in cancer cell viability, we utilized the AMPK inhibitor dorsomorphin and found that dorsomorphin treatment significantly decreased hepatoma cell viability (**Fig. 3F**). Additionally, both genetic knockdown (**Fig. 3G**) and pharmacological suppression of AMPK by dorsomorphin (**Fig. 3H**) substantially reduced hepatoma cell migration. These results underscore that AMPK is indispensable for IL6-enhanced hepatoma cell proliferation and migration.

To gain deeper insights into the mechanism through which AMPK exerts its regulatory influence on PFKL, we conducted experiments utilizing dorsomorphin and evaluated both PFKL protein and mRNA levels. Our observations revealed that treatment with the AMPK inhibitor led to a decrease in PFKL and pAMPK protein levels (**Fig. 3I**). Notably, as dorsomorphin also impacts TGFb-SMAD signaling, we performed western blot analyses for pAMAD5 and pACC1/2 in the dorsomorphin-treated cells and observed reductions in their levels as well (**Fig. 3I**). Intriguingly, the AMPK inhibitor had no discernible effect on PFKL mRNA levels (**Fig. 3J**). Additionally, we found that knockdown of AMPK resulted in an increase in proteasome activity in PLC5 cells (**Fig. 3K**). In summary, our findings collectively support the conclusion that AMPK stabilizes the PFKL protein through the inhibition of proteasome activity, unveiling a novel mechanism of posttranslational regulation involving AMPK, PFKL, and RPIA.

### IL6 increases the transcription of AMPK and PFKL via pSTAT3 directly binding to the promoters of AMPK and PFKL

Given that AMPK knockdown does not influence the IL6-mediated activation of pSTAT3 (**Fig. 3C**), it raises the possibility that pSTAT3 may function upstream of AMPK and PFKL in the regulatory cascade. To further explore this possibility, we applied two STAT3 inhibitors, nifuroxazide and BBI608, to investigate how STAT3 governs the expression of AMPK and PFKL. Employing these two STAT3 inhibitors effectively curtailed IL6-induced transcription of both AMPK and PFKL (**Fig. 4A and 4B**), providing compelling evidence that STAT3 modulates AMPK and PFKL expression. Through chromatin-immunoprecipitation assays utilizing pSTAT3 antibody and promoter fragments of AMPK and PFKL, we successfully demonstrated that pSTAT3 exerts its influence by directly binding to the promoters of both AMPK and PFKL (**Fig. 4C**). This direct interaction results in the augmented transcription of AMPK and PFKL, firmly establishing the AMPK and PFKL genes as direct transcriptional targets of pSTAT3 in this regulatory pathway.

**Figure 4.**
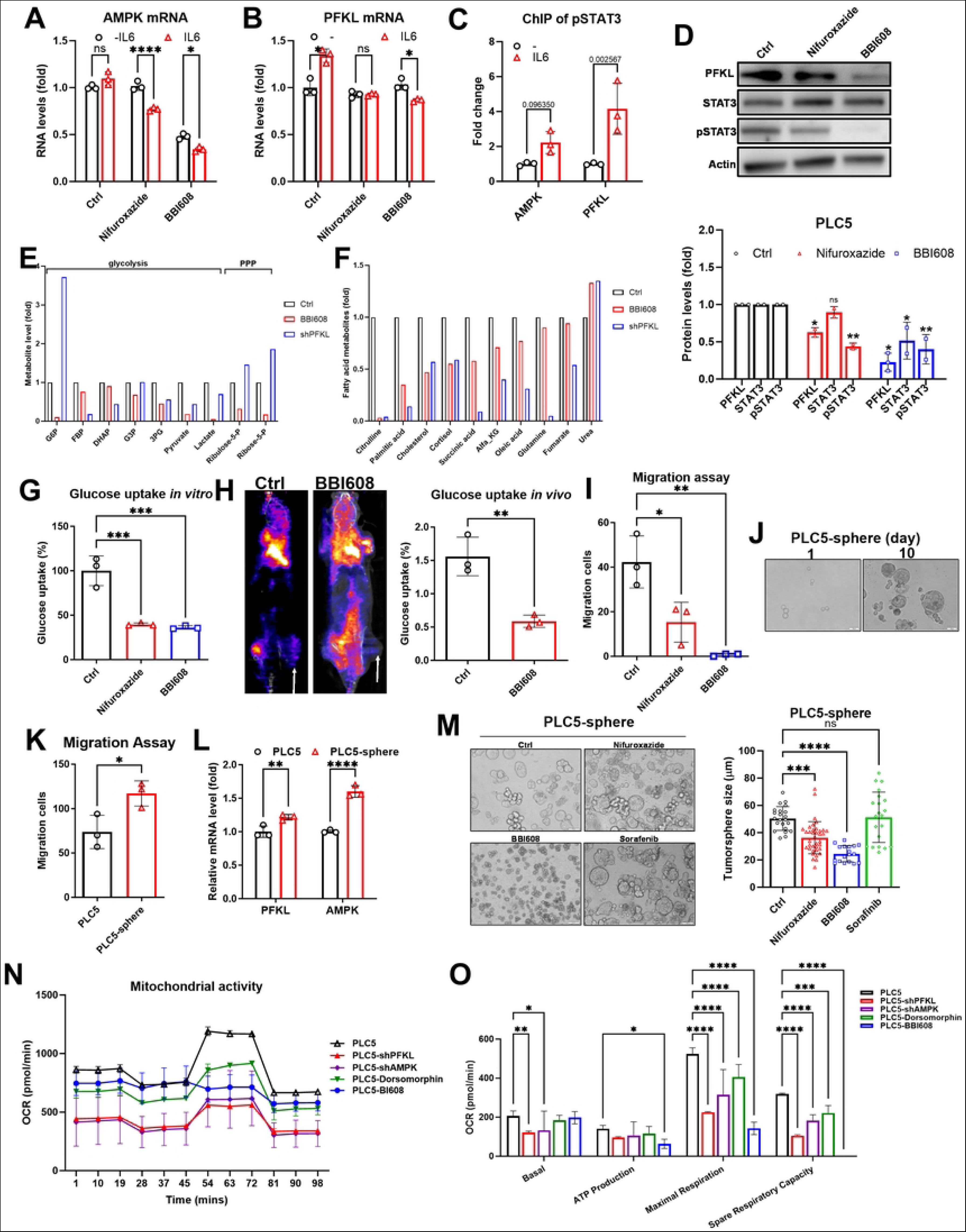
STAT3 directly regulates AMPK and PFKL transcription, resulting in the modulation of glycolysis in HCC. (**A**) Treatment with the STAT3 inhibitors nifuroxazide and BBI608 reduces AMPK mRNA levels in PLC5 cells treated with IL6. (**B**) Nifuroxazide and BBI608 treatments block IL6-induced PFKL-upregulated mRNA in PLC5 cells. (**C**) Chromatin immunoprecipitation assays reveal that pSTAT3 directly binds to the promoter regions of AMPK and PFKL in IL6-treated PLC5 cells. (**D**) Inhibition of STAT3 by nifuroxazide and BBI608 diminishes the IL6-mediated increase in PFKL protein levels, as shown by Western blot analysis. The lower panel quantifies the western blot results for PFKL, STAT3, and pSTAT3 protein levels. Red bars represent nifuroxazide treatment, blue bars denote BBI608 treatment, and black bars represent the no treatment control. (**E**) Treatment with the AMPK inhibitor BBI608 reduces the levels of glycolytic and lipogenic metabolites. Red bars represent the BBI608 treatment, while black bars represent the no treatment control. (**F**) Knockdown of PFKL with shPFKL leads to a significant reduction in glycolytic and lipogenic metabolites. Red bars indicate shPFKL, and black bars represent shLuc control. (**G**) Nifuroxazide and BBI608 treatments markedly decreased glucose uptake in PLC5 cells, as indicated by a reduction in (**H**) a tumor xenograft model, detected using FDG nuclear imaging. Arrows indicate the xenograft sites. Quantification of glucose uptake at the xenograft sites is shown. Red bars represent the BBI608 treatment, while black bars denote the no treatment control. (**I**) Targeting STAT3 with nifuroxazide and BBI608 reduces the migration ability of PLC5 cells. Red bars indicate nifuroxazide treatment, blue bars denote BBI608 treatment, and black bars represent the no treatment control. (**J**) PLC5 cells cultured in spheres as cancer stem cells for 10 days under specific conditions. (**K**) PLC5 tumorspheres exhibit increased migration ability compared to parental cells. (**L**) PLC5 tumorspheres display elevated mRNA expression levels of PFKL and AMPK. (**M**) The size of PLC5 tumorspheres can be reduced by STAT3 inhibitors but not by sorafenib. (**N**, **O**) Knockdown of PFKL and AMPK and suppression of AMPK and STAT3 resulted in a remarkable reduction in mitochondrial respiration and ATP production. Statistical analyses were performed using one-way ANOVA. *p < 0.05; **p < 0.01; ***p < 0.001; ****p < 0.0001.

### Targeting STAT3 to counteract glycolytic and mitochondrial activities in HCC

Treatment with two STAT3 inhibitors, nifuroxazide and BBI608, resulted in a significant reduction in PFKL protein levels under IL6 induction conditions (**Fig. 4D**). Furthermore, the suppression of STAT3 led to a notable decrease in glycolytic metabolites, including fructose 1,6-bisphosphate (FBP), dihydroxyacetone phosphate (DHAP), 3-phosphoglycerate (3PG), pyruvate, and lactate, in PLC5 cells, which paralleled the effects observed upon PFKL knockdown (**Fig. 4E**). Intriguingly, fatty acid metabolites such as cholesterol, oleic acid, palmitic acid, and succinic acid also exhibited reductions following STAT3 suppression or PFKL knockdown (**Fig. 4F**). Additionally, citrulline and cortisol levels were notably reduced (**Fig. 4F**). Suppression of STAT3 further translated to a decrease in glucose uptake both *in vitro* (**Fig. 4G**) and *in vivo* using the 18F-FDG nuclear imaging platform (**Fig. 4H**). Moreover, treatment with either nifuroxazide or BBI608 significantly diminished PLC5 cell migration *in vitro* (**Fig. 4I**), mirroring the effects of PFKL knockdown (**Fig. 1F**). Collectively, these findings highlight pSTAT3 as the principal mediator in conjunction with IL6 on the glycolytic rate, cell proliferation, and migration in HCC.

We next asked whether targeting STAT3 could be a viable therapeutic strategy against liver cancer. We induced tumorsphere formation in IL6/EGF/bFGF/HGF-treated PLC5 cells (**Fig. 4J**) and observed that PLC5-derived tumorspheres exhibited enhanced migration capacity (**Fig. 4K**) and higher expression of PFKL and AMPK (**Fig. 4L**). Notably, the size of PLC5-derived tumorspheres could be significantly reduced by the two STAT3 inhibitors nifuroxazide and BBI608 but not by the FDA-approved anti-HCC agent sorafenib (**Fig. 4M**). These results suggest that targeting STAT3 may represent a more effective approach against cancer stem-like cells.

Given the known connections of PFKL and AMPK to energy production, we proceeded to measure mitochondrial activity and ATP production via seahorse analysis following PFKL and AMPK knockdown, as well as AMPK suppression by dorsomorphin and STAT3 inhibition by BBI608. Our results indicated that all the aforementioned treatments led to a decrease in mitochondrial respiration in PLC5 cells (**Fig. 4N**). Importantly, the inhibition of STAT3 by BBI608 resulted in a remarkable reduction in ATP production (**Fig. 4O**), akin to the effects observed with PFKL knockdown (**Fig. 4O**). These findings collectively suggest that IL6 activates pSTAT3, which, in turn, binds to the promoters of AMPK and PFKL to enhance mRNA expression, glycolytic rate, and mitochondrial activity in HCC. Therefore, targeting STAT3 has emerged as a promising therapeutic approach against HCC.

### The PFKL positive glycolytic body is induced by hypoxia and low serum, and the direct PFKL-RPIA interaction explored through BiFC assay

PFKL stands out as a critical component in the glycolytic body (G-body), a central hub associated with tumorigenesis [18]. In our investigation, we noted that while the vector pAcGFP1-N2 exhibited a diffuse GFP signal in the cytoplasm, the PFKL-GFP fusion protein displayed distinctive aggregations within hepatoma cell lines (Hep3B, PLC5, HepG2, and Huh7) but remained diffuse in normal human liver cell lines (L02) (**Fig. 5A**). Notably, G-body formation alongside RNA [27], prompted experiments involving RNase A treatment, demonstrating large puncta formation in PLC5 cells that increased with IL6 treatment but became diffuse following RNAseA treatment (**Fig. 5B**).

**Figure 5.**
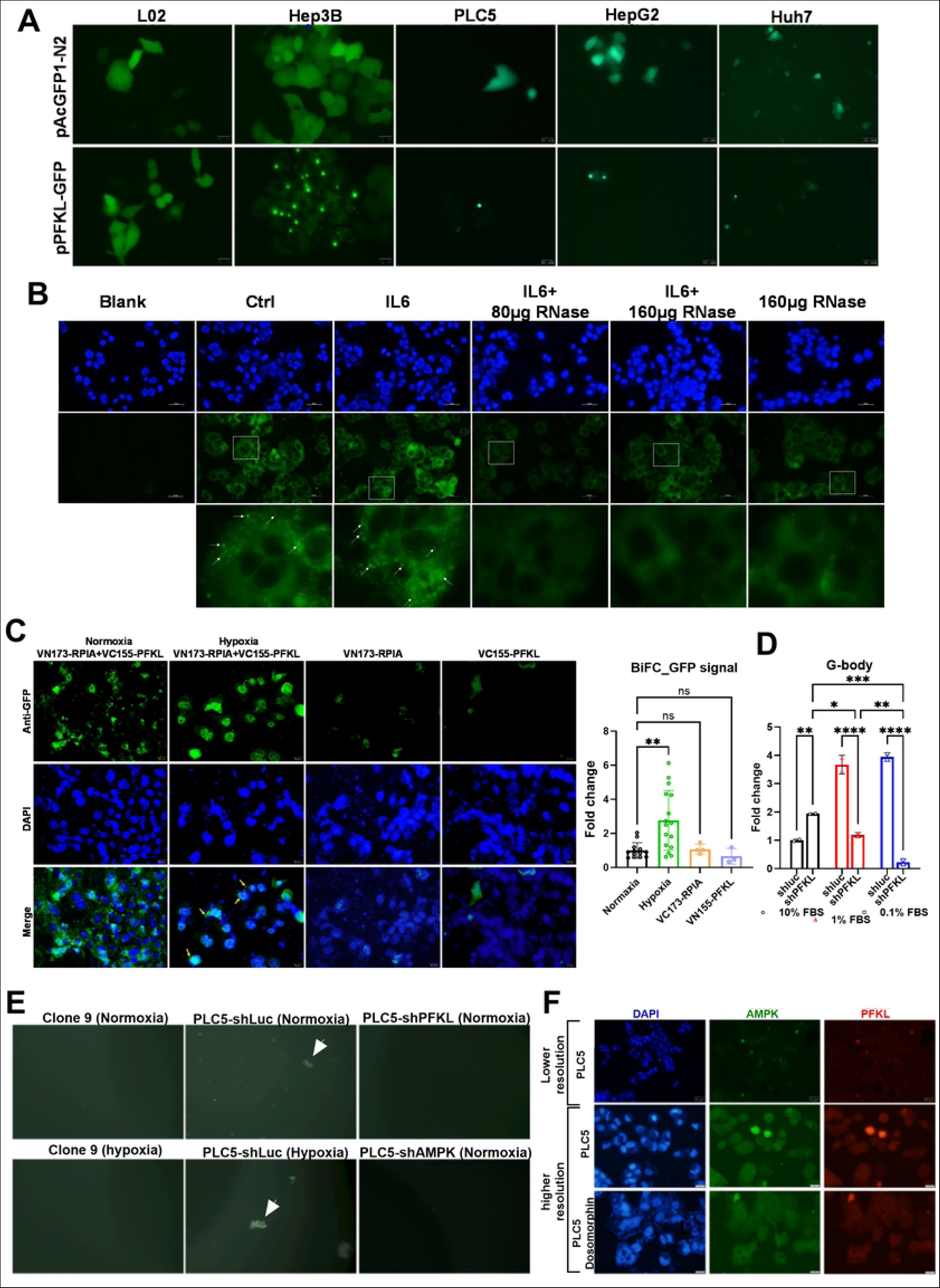
Unveiling G-Body Dynamics: PFKL’s Response to Hypoxia and Low Serum, and evidence of direct interaction between PFKL and RPIA using bimolecular fluorescence complementation (BiFC) assay. (**A**) PFKL-GFP forms clusters in hepatoma cells but not in normal liver cells. Fluorescence imaging shows PFKL-GFP in L02 normal hepatocytes and hepatoma cells (Hep3B, PLC5, HepG2, and Huh7). The upper panels display the GFP vector control, which exhibits a diffuse pattern of GFP in the cytoplasm, while the lower panels depict the cluster formation of PFKL-GFP in hepatoma cells but not in L02 cells. (**B**) Treatment with RNase reduces PFKL-assembled particles, as observed through immunostaining. IHC imaging demonstrates the PFKL protein pattern in PLC5 hepatoma cells after IL6 or RNase treatment, as well as IL6 plus RNase cotreatment. Arrows indicate the PFKL clusters that increase upon IL6 treatment and disappear after RNase cotreatment. (**C**) The BiFC assay provides evidence of the interaction between PFKL and RPIA. The upper panels show GFP signals in VN173-RPIA and VC-165 PFKL under normoxia, which diffuse in the cytoplasm but become aggregated in the cytoplasm or even enter the nucleus under hypoxia. GFP signals are absent in VN173-RPIA or VC155-PFKL alone. Scale bar: 100 µm. (**D**) Knockdown of PFKL specifically reduces the low serum induced G-body formation. (**E**) Hypoxia induces G-body formation in hepatoma cell (PLC5) but not normal liver cell (Clone 9), and knockdown of either PFKL or AMPK diminishes the G-body formation. (**F**) Co-localization of AMPK and PFKL can be abolished by suppression of AMPK with dorsomorphin. Statistical analyses were conducted using one-way ANOVA. ns: not significant; *p < 0.05; **p < 0.01; ***p < 0.001; ****p < 0.0001.

To substantiate the PFKL-RPIA interaction, a Bimolecular Fluorescence Complementation (BiFC) Assay was employed for live visualization. Fusing the nonfluorescent N-terminus to RPIA and the C-terminal fragment to PFKL, the expected lack of GFP production individually validated the interaction. Interestingly, the PFKL-RPIA interaction was observed in the cytoplasm under normoxic conditions but translocated to the nucleus during hypoxia Quantitative analysis of the BiFC signals revealed a significant increase, providing further evidence of the interaction (**Fig. 5C**). These in-depth analyses and accumulating evidence collectively contribute to a more comprehensive understanding of the direct interaction between PFKL and RPIA.

We further validated these findings by isolating G-bodies using Dynabeads, as previously described [18]. Low serum conditions promoted G-body formation in PLC5 cells, and the knockdown of PFKL substantially reduced G-body formation (**Fig. 5D**). Hypoxic stress similarly induced G-body formation in hepatoma cells but not in normal liver cells (Clone 9), with knockdown of PFKL or AMPK reducing G-body formation under these conditions (**Fig. 5E**).

Considering that AMPK can stabilize PFKL and that PFKL in turn stabilizes RPIA, we hypothesized that AMPK and RPIA may colocalize with PFKL within the G-body, acting as a crucial coordination center for glycolysis, the PPP, and carcinogenesis in hepatoma cells. Our investigation indeed revealed the colocalization of AMPK and PFKL through immunostaining, with the staining of both AMPK and PFKL being reduced upon treatment with the AMPK inhibitor dorsomorphin (**Fig. 5F**). These findings highlight the significance of PFKL and AMPK in G-body formation within hepatoma cells.

### Clinical relevance of the G-body: Colocalization of PFKL, RPIA, AMPK, and PKM2 within the G body in human liver cancer specimens

As our previous research reported that RPIA stabilizes β-catenin protein levels and activates downstream target genes during cancer formation [28], we speculated that RPIA may interact with β-catenin within the G-body, stabilizing β-catenin and promoting its translocation into the nucleus during carcinogenesis. By employing liver cancer arrays, we conducted immunostaining for PFKL and RPIA to explore the clinical relevance of the G-body. Remarkably, we observed that both PFKL and RPIA formed prominent puncta as early as HCC stage I, with a noteworthy observation being that these PFKL/RPIA clusters were situated within the nucleus (**Fig. 6A**). Utilizing double fluorescence immunostaining with anti-RPIA and anti-PFKL or anti-PKM2 and anti-AMPK antibodies in liver disease arrays (**Fig. 6B**), we found that PFKL staining formed small aggregates in chronic hepatitis, with a substantial increase in the number of PFKL and RPIA double-positive clusters observed in cases of chronic active hepatitis, persisting from cirrhosis through HCC stage IVA (**Fig. 6B**).

**Figure 6.**
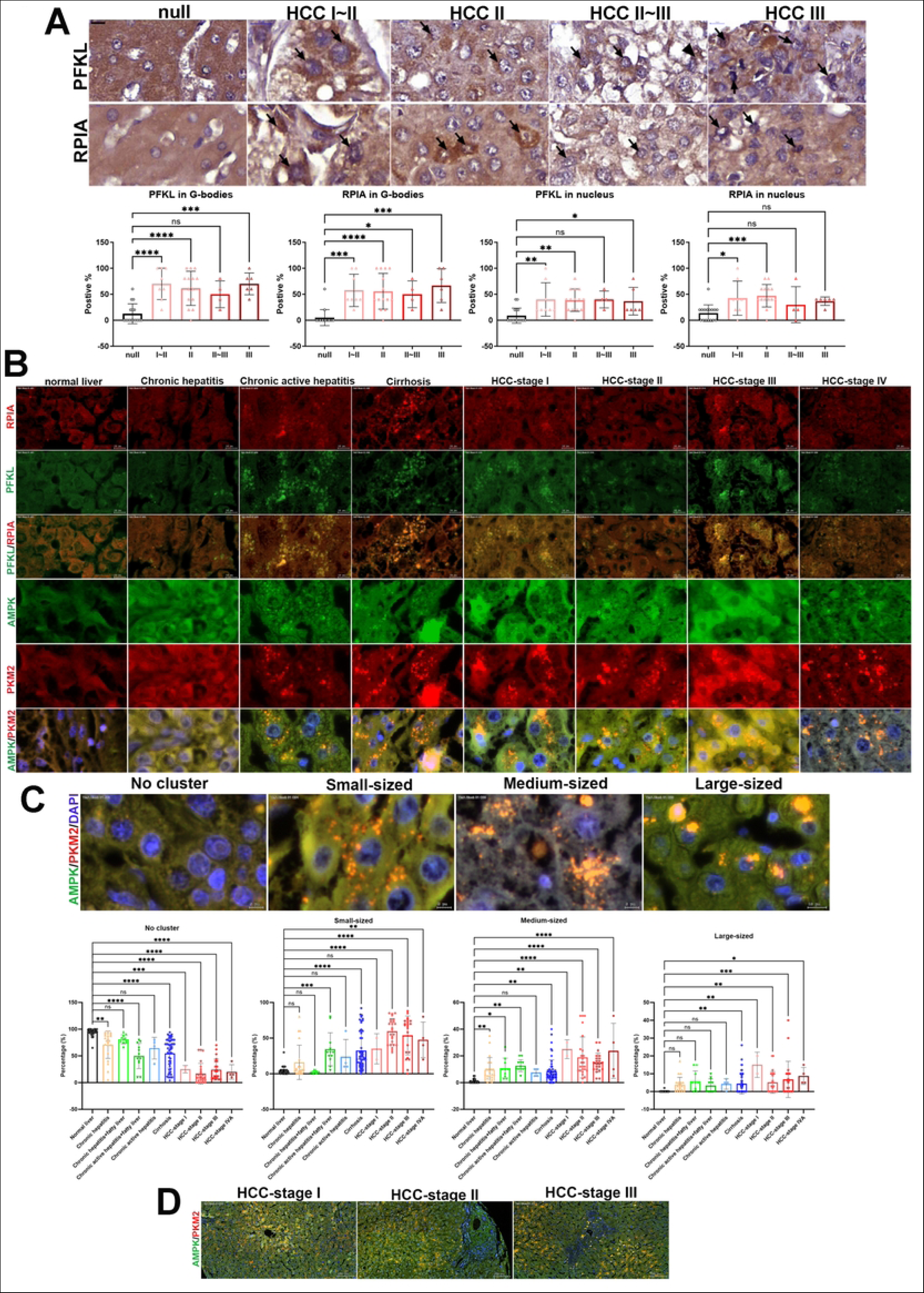
Aggregation of PFKL and RPIA in G-bodies in liver diseases and a cancer tissue array. (**A**) PFKL and RPIA from G-bodies in HCC specimens using a liver cancer tissue array. Arrows highlight the aggregation of PFKL- or RPIA-positive proteins. The lower panels present the quantification of immunohistochemistry results, demonstrating that the presence of PFKL and RPIA in G-bodies or the nucleus is positively correlated with HCC staging. (**B**) Double immunofluorescence staining reveals the colocalization of RPIA and PFKL, as well as AMPK and PKM2, in various stages of liver diseases, including chronic hepatitis, cirrhosis, and HCC, using a liver disease tissue array. The colocalization of RPIA/PFKL and AMPK/PKM2 is observed in small, medium, and large clusters, representing different stages of liver diseases. Scale: 10 µm. (**C**) Images depict the distribution of G-bodies in stages of liver disease, showing no cluster, small, medium, and large-sized clusters. The lower panels present quantification results for the presence of these clusters in different stages of liver disease. (**D**) Images illustrate the distribution of G-bodies in stage I, II, and III HCC specimens at lower magnification. Scale: 100 µm. Statistical analyses were conducted using one-way ANOVA. ns: not significant; *p < 0.05, **p < 0.01, ***p < 0.001.

A previous study indicated that medium-sized PFKL clusters are primarily responsible for diverting glucose flux into the pentose phosphate pathway in cells, whereas the larger-sized PFKL clusters tend to direct glucose flux toward serine biosynthesis in cancer cells [29]. Additionally, we investigated changes in G-body size within liver disease arrays via double fluorescence immunostaining with anti-AMPK and anti-PKM2 antibodies. We quantified the colocalization and statistical relevance based on clinical stages, which revealed an increase in the sizes and numbers of G-bodies. In cases of nodular cirrhosis, G-bodies displayed an increase in size and translocated toward the nucleus during hepatocarcinogenesis. In specimens of active hepatitis, small-sized clusters witnessed a dramatic increase (**Fig. 6C**). Intriguingly, we also observed that G-bodies were situated closer to the center of the liver lobe in cases of HCC stage I (**Fig. 6D**). From the stage of HCC onwards, the presence of medium- and large-sized clusters became strikingly evident, distributed away from the central vein, potentially conferring a hypoxic advantage to cancer cells. These findings derived from human liver specimens corroborate our discovery that the multienzyme metabolic complex G-body, consisting of PFKL, AMPK, RPIA, and PKM2, serves as a central hub for coordinating glucose metabolism, the PPP, and cancer formation in human liver diseases.

### Assessing the therapeutic efficacy of STAT3 and AMPK inhibitors in a zebrafish model

Subsequently, we investigated whether pharmacological targeting of STAT3/AMPK could serve as an effective therapeutic strategy for combating HCC *in vivo*. Employing a zebrafish model, we found that nifuroxazide notably curbed cancer cell proliferation in xenotransplantation assays (**Fig. 7A, A’**). Similarly, treatment with dorsomorphin, an AMPK inhibitor, also significantly reduced cancer cell proliferation in xenotransplantation (**Fig. 7A, A’**).

**Figure 7.**
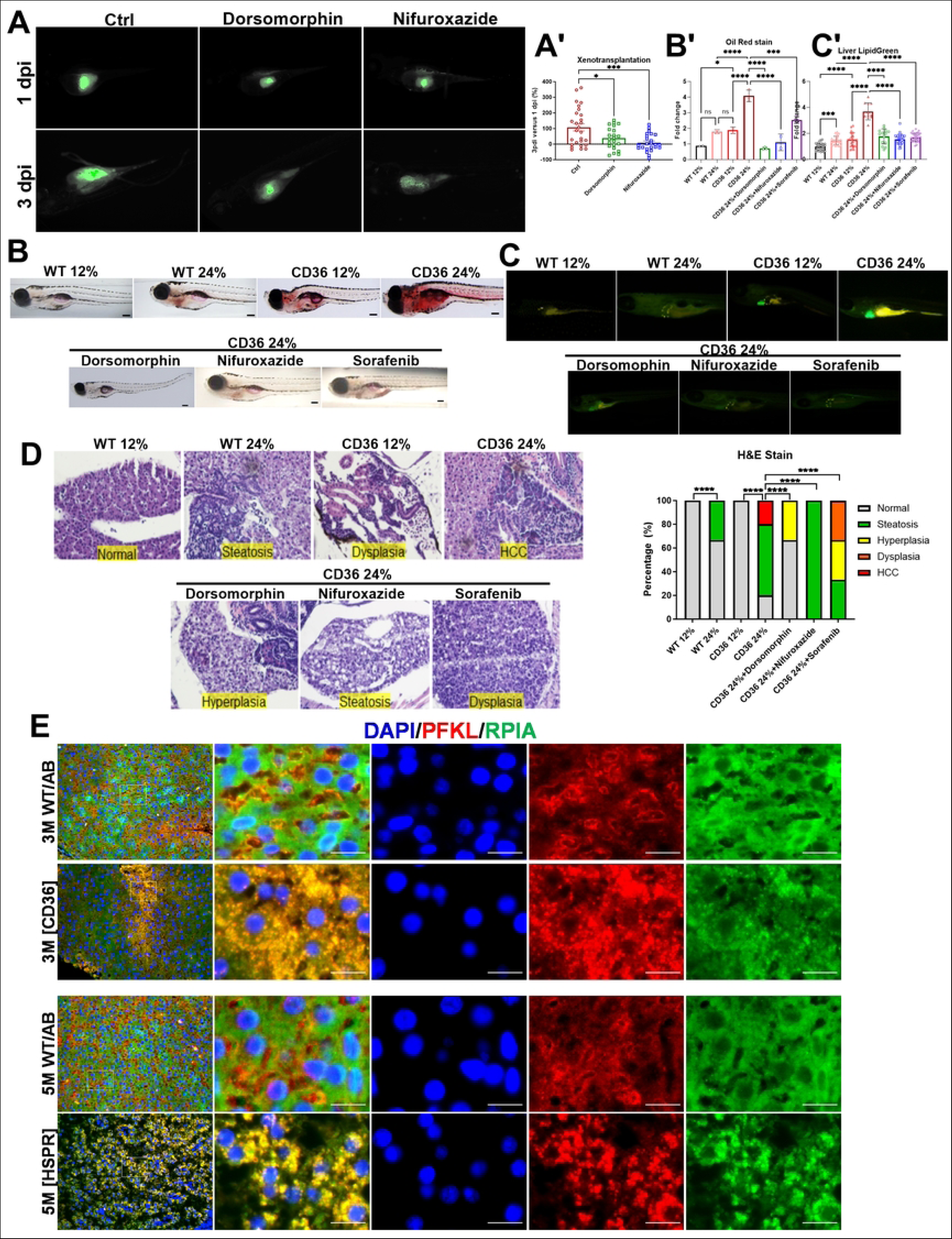
The therapeutic efficacy of AMPK and STAT3 inhibitors in zebrafish HCC models. (**A**) In a zebrafish xenograft model, suppression of STAT3 with nifuroxazide dramatically reduced HepG2 cell proliferation, while dorsomorphin also significantly decreased HepG2 cell proliferation, although to a lesser extent. Quantification results are presented in **A’**. (**B**) Oil red O staining demonstrates the anti-lipid accumulation effect of nifuroxazide and dorsomorphin in comparison to sorafenib in a 15-day-old CD36 transgenic fish model fed a high-fat diet. Quantification results are shown in **B’**. (**C**) LipidGreen2 staining yields similar results regarding the anti-lipid accumulation effect of nifuroxazide and dorsomorphin compared to sorafenib in the CD36 transgenic fish HCC model. Quantification results are presented in **C’**. (**D**) The anti-HCC effect of nifuroxazide and dorsomorphin was compared to that of sorafenib in a 1-month-old CD36 transgenic fish model fed a high-fat diet. Quantification results for H&E staining are shown in the right panel. Statistical analyses were performed using one-way ANOVA. *p < 0.05; **p < 0.01; ***p < 0.001; ****p < 0.0001. (**E**) Representative images of liver sections from 3-month-old wild-type (WT) and CD36 zebrafish and 5-month-old WT and HSPR (HBx, src, p53-/-, RPIA) zebrafish stained with the indicated antibodies. Images in the second column provide magnified views of the window area from the first column. Scale bars represent 10 μm.

We further demonstrated that both nifuroxazide and dorsomorphin displayed a more pronounced anti-lipid accumulation effect than sorafenib in the CD36 (fatty acid translocase) transgenic zebrafish model [30] under the high-fat diet by Oil Red O staining (**Fig. 7B, B’**). Consistent with those from Oil Red O staining, similar results were obtained by quantifying lipid accumulation in the livers of CD36 transgenic fish fed a high-fat diet with or without the different treatments by using Lipid Green2 staining (**Fig. 7C, C’**).

In our prior research, we utilized CD36 transgenic zebrafish exposed to a high-fat diet regimen for 30 days to induce HCC formation, which serves as a model akin to NASH-induced HCC [30]. In this study, we observed that nifuroxazide treatment can prevent the progression to HCC and maintain a steatotic state in CD36 transgenic zebrafish fed a high-fat diet (**Fig. 7D**). Furthermore, dorsomorphin treatment also blocked HCC formation and resulted in only some hyperplasia phenotype and an impressive amount of normal phenotype up to 65% in those fish populations (**Fig. 7D**). These results indicate that dorsomorphin and nifuroxazide exhibit potent anti-HCC properties in the CD36 transgenic zebrafish high-fat diet model. Notably, sorafenib exhibited lower efficacy in inhibiting the progression of HCC formation than dorsomorphin and nifuroxazide (**Fig. 7D**). Collectively, these findings indicate that targeting STAT3 and AMPK may offer promising therapeutic avenues for the treatment of liver cancer.

To extend the validity of the finding, we utilized two zebrafish HCC models to assess whether G-bodies could be detected in the zebrafish liver under HCC: one involving CD36 on a normal diet at 3 months and the other featuring HBx, src, RPIA, and p53-transgenic fish that developed HCC at 5 months. In both models, we consistently observed strong colocalization of PFKL and RPIA as aggregates, which were notably positioned closer to the nucleus (**Fig 7E**). These findings underscore the relevance of our observations in hepatoma cells and human liver tissues and support the presence of G-bodies in zebrafish liver cancer models.

## Discussion

In this investigation, we embarked on a comprehensive study of PFKL in tumorigenesis, shedding light on its multifaceted impact on tumor-related processes, encompassing glycolysis, ATP generation, cell proliferation, and migration. Our findings unveiled a pivotal role for PFKL in stabilizing RPIA protein levels through the ubiquitination/proteasome pathway, thereby establishing a crucial specific link between PFKL in glycolysis and RPIA in the pentose phosphate pathway. Dysregulation of PFKL and RPIA engenders significant alterations in glucose and lipid metabolism.

Furthermore, our study delved into the involvement of AMPK in hepatoma cell survival and migration, introducing the AMPK inhibitor dorsomorphin as a prospective therapeutic agent. Our data elucidated the significant regulatory role of AMPK in IL6-mediated PFKL expression in PLC5 cells. Remarkably, we also revealed that AMPK functions as a promoter, rather than a suppressor, in liver cancer by IL6-induced PFKL, a finding that challenges conventional perceptions. Although AMPK is known to suppress mTOR, which is typically regarded as a tumor suppressor [31, 32], it can simultaneously stimulate protumorigenic processes, including fatty acid oxidation and mitochondrial metabolism [24]. The theory, suggesting that AMPK activators have potential against HCC, is based on studies investigating metformin as an AMPK activator [33], but a recent study indicated that metformin also inhibits mTOR [34]. Combining metformin with an AMPK inhibitor might represent a more efficacious therapeutic strategy against HCC [24], although further investigations are warranted to elucidate the precise role of AMPK in HCC development. Notably, our study offers the pioneering insight that AMPK plays a role in stabilizing PFKL, a factor contributing to cell survival and migration in HCC.

Moreover, we provided compelling evidence that IL6 mediates HCC tumorigenesis via the activation of pSTAT3, which subsequently triggers AMPK and PFKL transcription. We demonstrated that targeting AMPK and STAT3 effectively curtails hepatoma cell proliferation and migration. Importantly, nifuroxazide emerged as a potent inhibitor of hepatoma cell lines while sparing normal liver cells, thus meriting consideration as a potential therapeutic agent against HCC (**Fig 8**).

**Figure 8.**
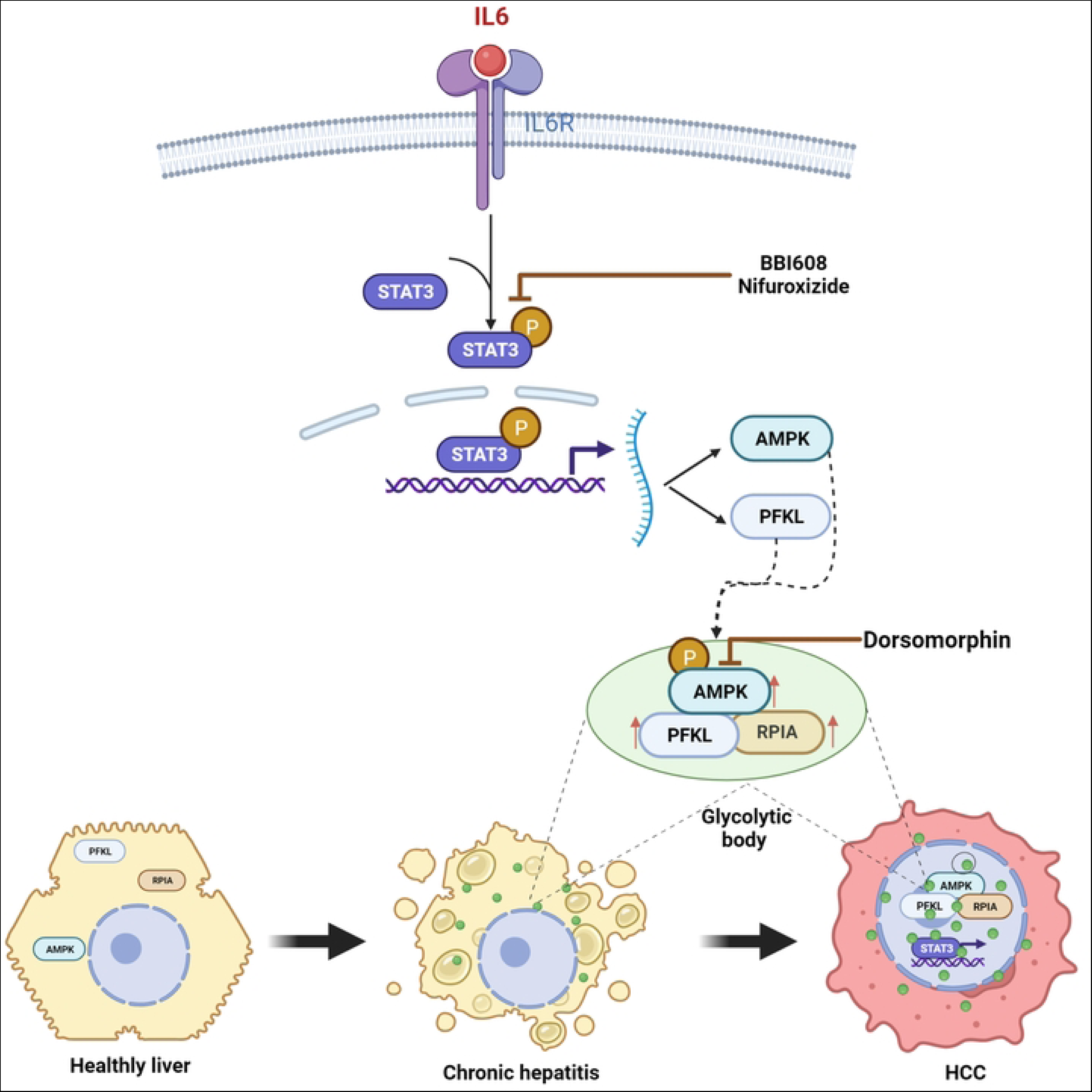
Model for IL6/STAT3-mediated AMPK/PFKL/RPIA stabilization in G-bodies during hepatocarcinogenesis. pSTAT3 increases the transcription of AMPK and PFKL. AMPK stabilizes the PFKL protein level. PFKL specifically stabilizes RPIA. AMPK, PFKL and RPIA are colocalized in the G-body. Inhibiting STAT3 or AMPK can reduce tumor proliferation and metastasis.

Notably, nifuroxazide, a STAT3 inhibitor bioactivated by ALDH1, confers tumor cell-specific action [35], explaining its favorable safety profile compared to BBI608 and sorafenib. Other STAT3 inhibitors, such as napabucasin (BBI608) [36, 37], exhibited broad inhibitory effects on both hepatoma and normal liver cells, underscoring the notion that generalized STAT3 inhibition might not constitute an ideal therapeutic strategy. Ideally, therapeutic targets for HCC should be tailored to HCC-specific gene expression, such as ALDH1 [38, 39] and IL6 [40]. IL6 trans-signaling through soluble IL6R is another potential target [41], in which targeting IL6Rα by miR218 and miR34a was demonstrated to diminish HCC [12]. Nevertheless, since IL6/STAT3 represents just one of many oncogenic signaling pathways in HCC, a more holistic approach combining STAT3 inhibition with other major pathway inhibitors may enhance therapeutic efficacy.

Numerous membraneless organelles emerge through liquid‒liquid phase separation in response to various stress conditions [42], which are linked to diseases such as Alzheimer’s disease and cancer [43, 44]. Targeting stress granules provides a promising avenue for cancer treatment [45]. The clustering of Wnt/β-catenin signaling also emerges as a potential novel target for cancer therapy [46]. The regulatory impact of ubiquitination on biomolecular condensate dynamics is implicated in various diseases, including cancer [47]. The pivotal role of PFKL in governing glycolytic rates is of paramount significance. Tumor cells prefer aerobic glycolysis, commonly known as the Warburg effect, to sustain their heightened proliferative rate [48]. Larger clusters of PFKL direct glucose flux [29], with PFKL compartmentalization at two interfaces (1 and 2), where interface 2 is particularly crucial for modulating associated protein assemblies with the cytoskeleton [49]. This clustering of PFKL plays a vital role in its biological function.

Our study uncovers the substantial influence of PFKL on liver cancer cell survival and migration. The intriguing observation that PFKL can form G-bodies and interact with various proteins, glycolytic enzymes, metabolites, and chaperones in both yeast and HCC cells under hypoxic conditions adds a new dimension to our understanding [18]. These interactions appear to enhance glucose consumption and ATP production under hypoxia, possibly by stabilizing interacting proteins, thus preventing their proteasomal degradation, as demonstrated in this study. Importantly, our findings suggest that PFKL not only stabilizes glycolytic proteins but also nonglycolytic proteins, such as RPIA and ERK, in HCC cells. Additionally, the size of G-bodies seems to correlate with clinical stages, with medium-sized G-bodies observed in chronic hepatitis, followed by larger G-bodies in cirrhosis and HCC specimens. Moreover, these clusters migrate toward the nucleus during cancer formation. Collectively, our study provides tantalizing insights into the role of the G-body as a central hub coordinating various metabolic pathways, including glycolysis, the pentose phosphate pathway, lipogenesis, and carcinogenesis in hepatoma cells. However, further investigations are warranted to elucidate the specific protein components that interact with PFKL within the G-body in liver cancer.

The coordination between glycolysis and the pentose phosphate pathway in cancer cells appears to hinge on posttranslational protein stabilization. PFKL, the rate-limiting glycolytic enzyme, stabilizes RPIA protein levels, a critical enzyme in the pentose phosphate pathway. IL6 has been shown to induce PFKL and activate pAMPK, thereby stabilizing PFKL protein levels. Additionally, IL6 stimulates pSTAT3 and upregulates AMPK and PFKL transcription. The colocalization of PFKL, AMPK, RPIA, and PKM2 within the G-body correlates with the progression from hepatitis to hepatocellular carcinoma. Targeting AMPK and STAT3 with specific inhibitors disrupts the IL6/STAT3-AMPK/PFKL pathway, resulting in reduced hepatoma cell viability and mitigated HCC formation. Our study proposes a potential therapeutic strategy against HCC.

## Conclusion

In summary, our study sheds light on the novel intricate interplay between PFKL and RPIA in liver cancer development. We unveil a regulatory mechanism wherein PFKL stabilizes RPIA protein levels through the ubiquitination-proteasome pathway, establishing a crucial link between glycolysis and the pentose phosphate pathway. Furthermore, our study delineates the multifaceted role of IL6 in HCC, demonstrating its capacity to induce PFKL expression via AMPK activation for protein stabilization and signal transducer and STAT3 activation for transcriptional upregulation. The colocalization of PFKL, AMPK, RPIA, and PKM2 within glycolytic bodies provides new insights into their coordination in regulating glucose metabolism and HCC progression. Importantly, inhibiting the IL6/STAT3-AMPK/PFKL axis emerges as a promising therapeutic strategy, as validated in zebrafish models. This comprehensive study of the molecular pathways governing metabolic dysregulation in liver cancer opens avenues for targeted therapeutic interventions against hepatocellular carcinoma.

## Materials and Methods

### Cell culture

Multiple human liver cancer cell lines, Hep3B (BCRC Cat# 60434, RRID:CVCL_0326), PLC5 (BCRC Cat# 60223, RRID:CVCL_0485), HepG2 (BCRC Cat# 60025, RRID:CVCL_0027), and Huh7 (CLS Cat# 300156/p7178_HuH7, RRID:CVCL_0336), were used to study the common mechanism. Hep3B cells have a p53 null mutation, PLC5 cells carry an R249S mutation at p53, whereas Huh7 cells have point mutations at p53 codon 220; these cell lines represent early (well-differentiated) HCC stages [50]. We also used mouse normal liver cell Clone 9 and normal human liver cell lines (L02) as controls. These cell lines were obtained from the Bioresource Collection and Research Center in Taiwan as previously described [20]. The normal liver Clone 9 cell line was a gift from the Institute of Nuclear Energy Research, Atomic Energy Council, Taoyuan, Taiwan.

The liver cancer cell lines HepG2, Huh7, Hep3B, and PLC5 were free of mycoplasma and cultured in Dulbecco’s modified Eagle’s medium (DMEM) supplemented with 10% fetal bovine serum (FBS), 100 units/ml penicillin, and 100 µg/ml streptomycin. The normal liver Clone 9 cell line was cultured in ham’s F12 medium supplemented with 10% FBS, 100 units/ml penicillin, and 100 µg/ml streptomycin. The cell lines were used in this study after being reauthenticated through short tandem repeat profiling (Applied Biosystems, Massachusetts, USA). All cells were incubated in a 37 °C incubator with 5% carbon dioxide.

### Quantitative polymerase chain reaction (qPCR)

The qPCR was conducted in accordance with a previously established protocol [20]. In brief, RNA extraction utilized the NucleoSpin® RNA Midi kit (MACHEREY-NAGEL, US), and cDNA synthesis was carried out using the iScriptTM cDNA synthesis kit (Bio-Rad, US) with a 1000 ng RNA template, following the manufacturer’s instructions. The qPCR procedure employed a SYBR Green system (Applied Biosystems, Foster City, CA, USA) with 20X diluted cDNA in a 384-well plate. The qPCR program included a hold stage at 95 °C for 20 seconds, a PCR stage at 95 °C for 1 second, and 60 °C for 20 seconds over 40 cycles, followed by a melt curve stage at 95 °C for 1 second and 60 °C for 20 seconds. To minimize technical errors, the qPCR analysis was performed in triplicate for each sample. The primer sequences are provided in Table S1.

### Gene knockdown and overexpression

Gene knockdown was conducted using small interfering RNA (siRNA) with Lipofectamine 2000 (Invitrogen) or a short-hairpin RNA (shRNA)-expression lentivirus system. The siPFKL containing the three Stealth RNA^TM^ siRNAs (HSS107868, HSS182257 and HSS182258, Invitrogen), the siRPIA containing three Stealth RNA^TM^ siRNAs (HSS117931, HSS117932 and HSS117933, Invitrogen), and the si-NC (negative control) containing the target sequence of 5’-UUCACUUCACUCCAUUUGUGUACC-3’, Invitrogen, Cat. 2935112). The specific shRNA (target sequence of PFKL: CTGAAGATGCTGGCACAATAC; AMPK: GTTGCCTACCATCTCATAATA) in the vector pLKO.1-puro was generated in 293T cells. The procedure was followed by our previous studies [20, 51]. PFKL and RPIA overexpression was performed using pCMV-PFKL (Origene, SC319353) and pcDNA3.0-RPIA, respectively, together with pcDNA 3.0 (Invitrogen) as the control.

### Western blots

Western blot analysis was carried out as described previously [20, 51]. The specific antibodies against PFKL (Cell Signaling Technology Cat# 8175, RRID:AB_11178807), STAT3 (Cell Signaling Technology Cat# 9132, RRID:AB_331588), pSTAT3 (Cell Signaling Technology Cat# 9130, RRID:AB_330367), AMPK (Cell Signaling Technology Cat# 2603, RRID:AB_490795), pAMPK (Cell Signaling Technology Cat# 5759, RRID:AB_10949320), ERK (GeneTex Cat# GTX59618, RRID:AB_10726211), pERK (Abcam Cat# ab32538, RRID:AB_11156273), RPIA (Abcam Cat# ab67080, RRID:AB_1142656), PK (GeneTex Cat# GTX111536, RRID:AB_1951258), G6P (GeneTex Cat# GTX113203, RRID:AB_2037119), TPI (GeneTex Cat# GTX104618, RRID:AB_1241405), pRaf (BioVision Cat# 3504-100, RRID:AB_2060496), pMEK1/2 (Cell Signaling Technology Cat# 9121, RRID:AB_331648), pSMAD5 (Abcam Cat# ab92698, RRID:AB_10561456), SMAD5 (Abcam Cat# ab40771, RRID:AB_777981), pACC (Cell Signaling Technology Cat# 3661, AB_330337), ACC (Cell Signaling Technology Cat# 3662, RRID:AB_2219400), pmTOR(Cell Signaling Technology Cat#2974, RRID: AB_2262884), mTOR (Cell Signaling Technology Cat#2983, RRID: AB_ 2105622), α/β-Tubulin (Cell Signaling Technology Cat#2148, RRID: AB_2288042), GAPDH (GeneTex Cat# GTX100118, RRID:AB_1080976), β-actin (GeneTex Cat# GTX109639, RRID:AB_1949572) and ubiquitin (Cell Signaling Technology Cat# 3936, RRID:AB_331292) were purchased from Cell Signaling (Danvers, Massachusetts, USA), Abcam (Cambridge, Massachusetts, USA), and GeneTex (Irvine, CA, USA).

### Cell viability assay

For the cell viability assay, we employed the MTT (3-(4,5-dimethylthiazol-2-yl)-2,5-diphenyltetrazolium bromide) or WST-1 (Water-Soluble Tetrazolium 1) assay from Takara, Japan, following the manufacturer’s protocol as previously outlined [28]. A 96-well plate was utilized for seeding 10^3^ × PLC5-shPFKL and PLC5-shLuc cells, with four replicates performed to assess cell viability in a time-dependent manner. The same experimental approach was adopted for evaluating the viability of PLC5-shAMPK and PLC5-shLuc cells, both with and without 20 ng/mL IL6 treatment. To examine the cytotoxic impact of the AMPK activator and inhibitor, PLC5 cells were subjected to dorsomorphin treatment in a dose-dependent manner for 48 hours.

### *In vitro* cell migration assay

A transwell migration assay (8 μm) as described earlier [28] was used to detect cell migration capacity in PLC5shPFKL or PLC5AMPK compared to PLC5shLuc, as well as measuring PLC5 cell migration treated with/without 6 μg/mL dorsomorphin. In brief, 5 × 10^4^ cells were placed in the upper layer of a cell culture insert with 200 μL of serum-free DMEM. Then, 750 μL of DMEM with 10% FBS and the test agent were loaded into a 24-well culture plate. Cells were incubated in a 37 °C incubator with 5% CO_2_ for 16 h. The membrane inserts were fixed in 3.7% formaldehyde for 5 min and subsequently incubated in 100% methanol for 20 min at room temperature. After 0.5% crystal violet in 2% ethanol to stain the membrane inserts for 15 min at room temperature, non-migrated cells on the upper membrane were scraped with cotton swabs. A PBS wash was carried out twice between operations. The cells that migrated through the membrane were imaged and counted using an inverted microscope (Axio Observer 3, Zeiss, Oberkochen, Germany).

### Bimolecular fluorescence complementation (BiFC) assay

We generated pBiFC-VN173-RPIA and pBiFC-VC155-PFKL plasmids and introduced the pBiFC-VN173-RPIA and pBiFC-VC155-PFKL plasmids separately into Hep3B and PLC5 cells using a transfection reagent, followed by a 36 hour incubation period. Afterward, we replaced the culture medium. We fixed the cells with 4% paraformaldehyde for 10-15 minutes at room temperature. Subsequently, we conducted an immunofluorescence staining procedure involving blocking, incubation with an anti-GFP primary antibody, and a secondary antibody. Each incubation step was followed by thorough PBS washes. The BiFC assay was then performed by mounting the cover glass slips onto glass slides using mounting medium and sealing the edges with nail polish. We employed fluorescence microscopy to excite the BiFC signal and captured images to assess the interaction between PFKL and RPIA. Data analysis involved examining the fluorescence images to determine the occurrence of protein interaction through GFP reconstitution.

### Mass spectrometry for measuring glycolytic metabolites

The metabolites were determined by chromatography-tandem mass spectrometry. In brief, 2 × 10^6^ PLC5-shPFKL, PLC5-shLuc, and PLC5 cells treated with 1 μg/mL nifuroxazide and BBI608 were collected. The metabolites were extracted using chilled 80% methanol for 30 min incubation. Each solution was then centrifuged at 12,000 rpm for 10 min at 4 °C. The supernatant was collected and concentrated using lyophilization. The pellets containing metabolites were resolved in distilled H_2_O and transferred to autosampler vials for liquid chromatography-tandem mass spectrometry (LC‒MS/MS, ACQUITY UPLC I-Class/Xevo TQ-S IVD, Waters, Massachusetts, USA) analysis. A series of calibration standards were prepared, along with samples to quantify metabolites.

### ATP production measurement by a Seahorse XF analyzer

The assessment of mitochondrial activity and ATP production was conducted using a Seahorse XF analyzer (Agilent, California, USA) along with the mitochondrial stress kit (Agilent, California, USA). This approach aimed to measure the oxygen consumption rate (OCR) in PLC5-shPFKL in comparison to PLC5-shLuc and in PLC5 cells treated with 1 μL of BBI608 for a duration of 2 hours. For the experimental setup, 2 × 104 cells were seeded in DMEM 24 hours prior to the analysis. Subsequent to treatment with or without BBI608, 675 μL of DMEM without sodium bicarbonate was introduced to the cells, and OCR was measured in accordance with the manufacturer’s protocol.

### Proteasome activity measurement

Proteasome activity was determined by a proteasome activity fluorometric assay kit (Biovision, California, USA) based on an AMC-tagged peptide substrate system to measure the proteolytic activity of PLC5-shAMPK compared to the PLC5-shLuc control. The experimental steps were followed according to the manufacturer’s protocol. In brief, 2 × 10^5^ cells were collected and homogenized with 100 μL of 0.5% NP-40 in distilled H_2_O. Each 25 μL sample was individually treated with 1 μL of MG132 (proteasome inhibitor) or assay buffer. Then, each sample was added to 1 μL of AMC-tagged peptide substrate for a 30 min incubation. The fluorescence was measured at Ex/Em = 350/440.

### Glucose uptake measurement

A glucose uptake fluorometric assay kit (Biovision, California, USA) was used to measure glucose uptake according to the manufacturer’s protocol with PLC5 cells treated with 1 μg/mL nifuroxazide and BBI608 for 2 h. In brief, 2 × 10^3^ cells were seeded for two days. The cells were then replaced with 100 μL of serum-free DMEM for 24 h. After washing with PBS three times, 90 μL of KRPH buffer (20 mM HEPES, 5 mM KH_2_PO_4_, 1 mM MgSO_4_, 1 mM CaCl_2_, 136 mM NaCl, 4.7 mM KCl, pH 7.4) with 2% BSA was added for 40 min. The cells were treated with 1 μg/mL of nifuroxazide and BBI608 for 2 h and subsequently with 10 μL of 10 mM 2-DG for 20 min. After washing with PBS three times, 90 μL of extraction buffer was added, and the cells were frozen/thawed once, followed by heating to 85 °C for 40 min. The samples were kept at 4 °C for 5 min and then added to 10 μL of neutralization buffer. Each 50 μL sample was mixed with 50 μL of reaction mix (1 μL of PicoProb, 1 μL of enzyme mix in 48 μL of assay buffer) for 40 min at 37 °C. The fluorescence was measured at Ex/Em = 535/587.

### *In vivo* glucose uptake by FDG nuclear imaging

*In vivo* glucose uptake was determined by FDG nuclear imaging. A small animal PET/CT scanner (nanoScan PET/CT, Mediso, Massachusetts, USA) was used to measure glucose uptake in BNL tumor-bearing BALB/c mice. The mice were purchased from BioLASCO, Taiwan, and maintained under a 12 h light/dark cycle at 22 °C. Animal studies were approved by the Institutional Ethical Review Committee at the Institute of Nuclear Energy Research and were performed according to NIH guidelines on the care and welfare of laboratory animals. A total of 2 × 10^6^ tumor cells were subcutaneously injected into the hind leg and grown for 14 days to a tumor size of 300 mm^3^. One day prior to nuclear imaging, 10 mg/kg BBI608 was injected into the tail vein of the mice (n = 3 vs PBS n = 3). ^18^F-FDG (150 μCi) was injected into the tail vein, and radioactive imaging was acquired after 30 min. The PET scan was conducted for 15 min on the mice, followed by a 5 min CT scan. The radioactive signal value was measured as the percentage of injected radioactivity dose/gram (% ID/g).

### Chromatin immunoprecipitation (ChIP) assay

To assess the binding of pSTAT3 to the genomic regulatory regions of AMPK and PFKL, we utilized the ChIP assay. Specifically, 5 μg of the p-STAT3 antibody (catalog# 9131, Cell Signaling, Danvers, Massachusetts, USA) was introduced in the ChIP procedure using protein G agarose beads, as per the manufacturer’s protocol (Merck EZChIP kit). Subsequently, 2 microliters of ChIP samples were employed for qPCR analysis, following the established methodology outlined in previous studies [30, 52]. The qPCR analysis utilized gene-specific primers, the sequences of which are detailed in Table S1.

### RNaseA treatment and measurement of G-body formation

The following experiment was conducted using PLC5 cells. Briefly, a total of 2 × 10^4^ PLC5 cells were cultured in a chamber slide and treated with 20 ng/mL IL-6, either alone or in combination with 80 μg or 160 μg of RNase, for a duration of 24 hours. Subsequently, the cells were fixed with 4% paraformaldehyde in PBS for 15 minutes at room temperature. After two washes with PBS, the cells were subjected to a 0.2% Triton-X100 treatment in PBS buffer for 10 minutes at room temperature, followed by replacement with 2% BSA in PBS for 1 hour. Next, the primary antibodies targeting PFKL, at a concentration of 2 μg/mL, were added to the cells and allowed to incubate for 24 hours at 4 °C. Following three washes with PBS, the cells were treated with a secondary antibody, Alexa Fluor 488 goat anti-rabbit IgG, for 1 hour at room temperature. Finally, the cells on the slides were mounted using a Prolong Gold anti-fade reagent (Invitrogen, Massachusetts, USA), and individual slides were examined using an inverted microscope (Axio Observer 3, Zeiss, Oberkochen, Germany).

### Immunostaining for cell culture

Immunostaining was used to determine the protein levels as described earlier [28]. A total of 2 × 10^4^ cells were cultured in an 8-well chamber slide (Merck Millipore, Massachusetts, USA) and then fixed with 4% paraformaldehyde in PBS buffer for 15 min at room temperature. The cells were washed twice with PBS before being treated with 0.2% Triton X-100 in PBS buffer for 10 min at room temperature, which was subsequently replaced with 2% BSA in PBS for 1 h. The primary antibody was added and incubated at 4 °C overnight. After three washes with PBS, the samples were incubated with Alexa Fluor 488 goat anti-mouse IgG or goat anti-rabbit IgG and Alexa Fluor 546 goat anti-rabbit IgG secondary antibodies for 2 h at RT. Cells on the slides were mounted with a Prolong Gold anti-fade reagent with DAPI (Invitrogen, Massachusetts, USA), and then the individual slides were detected using an inverted microscope (Axio Observer 3, Zeiss, Oberkochen, Germany).

### Immunohistochemistry (IHC) and immunofluorescence (IF) staining for tissue array

A CSA3 Human Liver cancer-metastasis-normal tissue array (Super Biochip) was used for PFKL immunostaining. LVC481 Liver cancer and normal tissue array (Biomax) was used for PFKL and RPIA immunostaining. The LV20812b Liver Disease Spectrum Tissue Array (Biomax) was used for PFKL and RPIA, as well as AMPK and PKM2 immunofluorescence staining.

The tissue array slide was baked for 30 minutes at 60 °C to prevent tissue detachment from the slide. The tissue array was dewaxed by using nonxylene and ethanol. Sodium citrate (10 mM, pH 6.0) plus 0.05% Tween 20 was used for antigen retrieval at 95 °C for 20 min, and the samples were cooled at room temperature. For immunostaining, 3% H_2_O_2_/methanol was used to remove endogenous catalase. 5% goat serum blocking 1 hr, primary antibody (1:100) incubation 4℃ overnight, biotin-conjugated secondary antibody (Vector) 30 min, ABC reagent 30 min for amplifying the signal, DAB detection, Hematoxylin counterstain, and dehydrated by Nonxylene and ethanol, and mounting slide. For immunofluorescence staining, the sections were washed with PBST, blocked with BlockPRO™ 1 Min Protein-Free Blocking Buffer for 1 hr, and incubated with primary antibody (1:200) at 4 °C overnight. PBST washes were then performed with the following secondary antibodies: anti-AMPK mouse antibody (NBP2-22127SS, Novus Biologicals, 1:200), anti-PKM2 rabbit mAb (D78A4, Cell Signaling Technology, 1:200), goat anti-mouse Alexa Fluor® 488 (A28175, Thermo Fisher Scientific Inc., 1:500), and goat anti-rabbit Alexa Fluor® 546 (A11010, Thermo Fisher Scientific Inc., 1:500) for the double staining of AMPK and PKM2. The secondary antibodies were as follows: anti-PFKL mouse monoclonal antibody (sc-393713, Santa Cruz Biotechnology, 1:200), anti-RPIA rabbit polyclonal antibody (GTX66545, GeneTex, Inc., 1:200), goat anti-mouse Alexa Fluor® 488 (A28175, Thermo Fisher Scientific Inc., 1:2000), and goat anti-rabbit Alexa Fluor® 546 (A11010, Thermo Fisher Scientific Inc., 1:1000) for the double staining of PFKL and RPIA. For immunofluorescence staining, after overnight primary antibody incubation, secondary antibody incubation was performed at room temperature for one hour. DAPI reagent was used to stain the nucleus.

### Zebrafish husbandry

Zebrafish husbandry was conducted at the Taiwan Zebrafish Core Facility (TZCF), accredited by the Association for Assessment and Accreditation of Laboratory Animal Care International (AAALAC) since 2015. All protocols adhered to guidelines and regulations, with approval from the Ethics Committee: Institutional Animal Care and Use Committee (IACUC) of the National Health Research Institutes (reference NHRI-IACUC-109031-M1-A).

Zebrafish husbandry, embryonic toxicity and xenotransplantation assays were performed as previously described [53–55]. Wild-type and CD36 transgenic zebrafish were utilized, and embryos were collected after cross-mating, then incubated in E3 solution at 28 °C. Embryo surfaces were cleaned with 6% and 8% bleach solutions at 16-22 hours post-fertilization (hpf). Larvae were subjected to different diets (normal diet with 12% fat or high-fat diet (HFD) with 24% fat) during the experimental period. Transgenic embryos were separated into control and treatment groups, with 50 larvae each, and were fed from 5 days post-fertilization (dpf). Diets were administered four times daily, accompanied by 20 ml of paramecium as an additional food source for larvae incapable of feeding on the normal or HFD.

CD36 transgenic larvae were used in drug screening for two experiment durations: a 15-day experiment focusing on liver lipid accumulation inhibition and a 30-day experiment concentrating on liver cancer inhibition. In the 15-day experiment, larvae were fed from 5 dpf to 15 dpf, followed by a two-day fasting period until 17 dpf for clearer staining observations. Sacrifice occurred on 18 dpf. For the 30-day experiment, larvae were fed from 5 dpf to 30 dpf, followed by a two-day fasting period until 32 dpf, with sacrifice on 33 dpf. All experimental procedures were meticulously conducted in line with ethical standards.

### Drug treatment

Three different drugs were administered to zebrafish in distinct manners for two distinct objectives: inhibiting lipid accumulation and inhibiting liver cancer. Following the last feeding session of the day, a group of 50 larvae was transferred to a 9 cm Petri dish filled with water containing dissolved drugs or chemical compounds at specific concentrations. Subsequently, the larvae were incubated overnight. Sorafenib (0.1 µM), nifuroxazide (10 µM), and dorsomorphin (10 µM) were dissolved in dimethyl sulfoxide (DMSO), and treatment was initiated after the last feeding on 15 dpf until 17 dpf for the 15-day-old larvae. For the 30-day-old larvae, treatment was administered from 25 dpf until the conclusion of the fasting period. The concentrations were determined based on the results obtained from the embryo toxicity test.

### Oil red O staining

For Oil Red O staining, fifteen larvae from each 15-day-old group were selected and subjected to the following procedure. Initially, they were fixed overnight at 4 °C in a 4% paraformaldehyde solution. On the subsequent day, larvae were rinsed twice with phosphate-buffered saline (PBS) and then treated with 80% and 100% 1,2-propylene glycol for 20 minutes at room temperature. Subsequently, the larvae were stained in the dark overnight with 0.5% Oil Red O prepared with 100% propylene glycol. On the third day, larvae underwent two PBS washes, followed by treatment with 80% and 100% 1,2-propylene glycol for 20 minutes at room temperature to eliminate excess staining. Before microscopic observation and imaging, larvae were stored in 80% 1,2-propylene glycol. This staining protocol closely resembled that used in a previous study [21].

Following staining, the larvae were rinsed with 1x PBS and immersed in 300 μl of 4% NP-40 prepared with 100% isopropanol, and incubated at room temperature overnight. On the fourth day, 95 μl of the immersion was transferred into a 96-well plate, and the absorbance was measured at 490 nm and 570 nm for the quantification of lipid accumulation.

### LipidGreen2 staining

For LipidGreen2 staining, groups of 15-day zebrafish larvae underwent an overnight fasting period following their feeding. LipidGreen2, a small molecule probe for lipid imaging known for selectively staining neutral lipids in cells and fat deposits in live zebrafish [56], was utilized. In a 9 cm Petri dish, larvae were incubated in a 10 μM LipidGreen2 solution for 30 minutes. Subsequently, the Petri dish was replaced with water, and larvae underwent a 30-minute incubation for destaining. The larvae were then placed on agar, anesthetized with tricaine, and subjected to microscopic observation and imaging of liver fluorescence. The intensity of liver fluorescence in the larvae was quantified using ImageJ.

### Hematoxylin & eosin (H&E) staining

To perform Hematoxylin & Eosin (H&E) staining for HCC histopathological observation, fifteen larvae from each group of 30-day-old larvae were collected. The tissues designated for histopathological analysis were fixed using a 10% formalin solution, embedded in paraffin, sectioned at a thickness of 5 μm, and mounted on poly-L-lysine-coated slides. The sections were then stained with H&E. Histopathological characteristics of the larvae were examined under magnifications of 50x and 400x for detailed observation and analysis.

### Xenotransplantation assay in zebrafish

For dechorionation, 1 dpf zebrafish eggs were put into 0.003% PTU/E3 medium approximately 20 hpf, followed by 20 µg/ml pronase (ChemCruz, Santa Cruz Biotechnology, Texas, US) to break the membrane. The embryos were stirred and transferred to another glass beaker until completely dechorionated. For cell injection, approximately 9 x 10^5^ HepG2 cells were collected in PBS, and then 5 µl of CFSE (Life Technologies, Invitrogen, Massachusetts, US) was added to the cell suspension and incubated at 37 °C for 15 minutes. After being centrifuged and washed with PBS, the pellet was resuspended in 20 µl of PBS and kept at 37 °C. Next, 0.016% tricaine in PTU/E3 medium was used to anesthetize 2-day postfertilization fish. The prepared HepG2 cells labeled with CFSE were injected into the zebrafish embryo yolks via microinjection. The fish were maintained in PTU/E3 medium and put in an incubator gradually heated from 28 °C to 37 °C. Healthy and injected embryos at 3 dpf were collected and transferred into a 96-well plate under a fluorescence microscope. Fluorescent signals were taken using an ImageXpress® Micro device (Molecular Device, California, USA) at 3 dpf (1 dpi (days-post-injection) and 5 dpf (3 dpi). Quantification of the area of fluorescence change was calculated using MetaXpress 2.3 (Molecular Device, California, USA). Nifuroxazide and dorsomorphin (MedChem Express, New Jersey, USA) were added at 3 dpf and changed at 4 dpf.

### Immunofluorescence staining of liver-section slides

Slides were dewaxed and rehydrated with nonxylene and serial concentrations of ethanol (100%, 95%, and 70%), and then antigens were retrieved by 10 mM sodium citrate buffer with 0.05% Tween 20 (pH 6.0) at 95 °C for 10 minutes. After cooling at room temperature, the slides were washed with PBST one time. Next, slides were blocked with 5% goat serum at room temperature for 1 hour with 60 rpm shaking and stained with 1% goat serum containing antibodies against PFKL (GTX105697) and RPIA (sc515328) at 4 °C overnight with 60 rpm shaking, followed by washing with PBST three times. After being washed, slides were further stained with 2% goat serum containing Alexa Fluor goat-anti-rabbit 546, Alexa Fluor goat-anti-mouse 488, and DAPI at room temperature for 1 hour with 60 rpm shaking. Finally, the slides were washed with PBST three times and mounted with aqueous medium. Images were acquired by a Leica DMIRB inverted microscope equipped with an Olympus color CCD DP73 and objective lenses of 40X (0.75 N.A).

### Statistical analysis

Statistical analyses were conducted using GraphPad Prism V10 (GraphPad Software, Inc., California, USA). The specific statistical tests employed are described in the figure legends and included either a two-tailed Student’s t test or one-way ANOVA. The significance levels are indicated by asterisks as follows: ns for not significant, * for 0.01 < P ≤ 0.05, ** for 0.001 < P ≤ 0.01, *** for 0.0001 < P ≤ 0.001, and **** for P ≤ 0.0001.

## Supplementary Material

**Table S1.** Primer sequences for qPCR and ChIP

**Figure S1.** Confirmation of viability assays through FACS-based analysis.

**Figure S2.** Knockdown of PFKL does not affect the expression of other enzymes in glycolysis and PPP, and PFKL regulates RPIA and MYC protein expression.

## List of abbreviations

PFKL: 6-phosphofructokinase liver type
RPIA: Ribose 5-Phosphate Isomerase A
PPP: pentose phosphate pathway
IL6: interleukin 6
G body: Glycolytic body
BiFC: Bimolecular fluorescence complementation assay
HCC: Hepatocellular carcinoma
MALFD: Metabolic dysfunction-associated fatty liver disease
NAFLD: Nonalcoholic fatty liver disease
NASH: Nonalcoholic steatohepatitis
PFK1: Phosphofructokinase-1
STAT3: Signal transducer and activator of transcription 3
PFKFB3: 6-phosphofructo-2-kinase/fructose-2,6-biphosphatase 3
ERK: Extracellular signal-regulated kinase
AMPK: AMP-activated protein kinase
mTOR: target of rapamycin
NHRI: National Health Research Institutes
IRS: immunoreactivity scores
FACS: fluorescence-activated cell sorting
pERK: phosphorylated extracellular signal-regulated kinase
PK: pyruvate kinase
PGI: phosphoglucose isomerase
TPI: triose-phosphate isomerase
G6PD: glucose-6-phosphate dehydrogenase
TALDO: transaldolase
TKL: transketolase
CSCs: cancer stem cells
FBP: fructose 1,6-bisphosphate
DHAP: dihydroxyacetone phosphate
3PG: 3-phosphoglycerate
cDNA: Complementary DNA
qPCR: Quantitative polymerase chain reaction
siRNA: small interfering RNA
shRNA: short-hairpin RNA
DMEM: Dulbecco’s modified Eagle’s medium
FBS: fetal bovine serum
MTT: 3-(4,5-dimethylthiazol-2-yl)-2,5-diphenyltetrazolium bromide
WST-1: Water-Soluble Tetrazolium 1
OCR: oxygen consumption rate
ChIP: Chromatin immunoprecipitation
IHC: Immunohistochemistry
IF: immunofluorescence
TZCF: Taiwan Zebrafish Core Facility
IACUC: Institutional Animal Care and Use Committee
AAALAC: Association for Assessment and Accreditation of Laboratory Animal Care International
hpf: hours post-fertilization
HFD: high-fat diet
dpf: days post fertilization
DMSO: Dimethyl sulfoxide
PBS: phosphate-buffered saline
H&E: Hematoxylin & eosin staining
dpi: day post-injection

## Author contributions

H.-Y. H and C.-C. C. acquisition, analysis and interpretation of data; Y.-T. C. acquisition and analysis of data; B. P. S. technical support; C.-L. H. acquisition, analysis and interpretation of data; Y.-W. W. technical support; C.-C. K. intellectual contribution, discussion and metabolomics analysis; W.-C.W. intellectual contribution, discussion and tissue array analysis; J.-Y. W. acquisition and analysis of data; W.-C. S. acquisition and analysis of data; H.-K. L. technical support; W.-Y.Y. performed human liver cancer tissue array IHC; Y.-H.L. performed the zebrafish IHC experiment; K.-H. G. acquisition and analysis of data; D. W. L. acquisition and analysis of data; H.-D. W. provided the intellectual contribution, concept, design, and discussion and edited the manuscript; C.-H. Y. conceptualized and designed the study, supervised the study and wrote the manuscript.

## Data availability

All data generated or analyzed during the current study are included in this published article (and its supplementary information files).

## Declarations

### Ethics approval and consent to participate

Zebrafish experiments were approved by the Institution of Animal Care and Use Committee (IACUC) of the NHRI (protocol No. NHRI-IACUC-109031-M1-A).

